# All You Need Is Water: Converging Ligand Binding Simulations with Hydration Collective Variables

**DOI:** 10.1101/2025.06.27.661893

**Authors:** Marc Schulze, Tetiana Khakhula, Nicola Piasentin, Simone Aureli, Valerio Rizzi, Francesco Luigi Gervasio

## Abstract

Selecting appropriate collective variables (CVs) is a crucial bottleneck in enhanced sampling molecular dynamics (MD) simulations. Although progress has been made with data-driven and intuition-based approaches, optimal CVs remain system-specific. Meanwhile, simple geometric descriptors are still widely used due to their transferability. A promising, yet under-explored, candidate for a more efficient CV is solvation. Indeed, despite its central role in ligand binding and folding, the complexity of solvent behavior has hindered its widespread use. Here, we introduce a data-driven and automatic strategy to construct robust solvation-based CVs. Our method identifies critical hydration sites by analyzing the radial distribution function of water around a ligand. Remarkably, using only these hydration CVs within on-the-fly probability enhanced sampling (OPES) simulations, we successfully converge the binding free energy landscapes for a series of host-guest systems. These landscapes show excellent agreement with those from more computationally expensive benchmark methods. We further demonstrate that the choice of where to bias water is key to efficient convergence, providing clear guidelines for implementation. This work not only underscores the central role of water in molecular recognition but also offers a powerful and generalizable framework for enhancing the sampling of complex biomolecular events.

## I. INTRODUCTION

Water is a vital component of life^1^. As the most abundant molecule on Earth’s surface and in living organisms, it is involved in countless chemical and biological processes, such as protein folding and ligand binding. Rather than being a passive stage for these events, water actively orchestrates them. However, this pivotal role is often overlooked in favor of the solute remaining in the limelight. Adopting a water-centric perspective is essential for understanding these processes, yet this remains a significant challenge due to the complex dynamics of solvation being notoriously difficult to capture.

This complexity arises from water’s ability to act as both a hydrogen bond donor and acceptor, forming a fluctuating network of three-dimensional hydrogen bonds with itself and with solutes^2,3^. A static picture, such as that emerging from high-resolution X-ray crystallography which only resolves high-occupancy water oxygen atoms^4^, is insufficient to fully capture the dynamic nature of solvation. In the case of ligand binding, the change in crystallographic water sites in studies of congeneric ligands with similar binding poses but different affinities^5–8^ has drawn attention to the necessity of considering all relevant water molecules^9^. Water molecules that interact directly with the solute are defined as the hydration shell, while those surrounded by an unperturbed sea of other water molecules are considered bulk water. The constant movement of water molecules fuels a dynamic exchange between these populations.

A more quantitative view comes from thermodynamics. Solvation is not uniform. Instead, it reflects the surface chemistry, topology, and molecular size of the solute, giving rise to complex thermodynamic landscapes^10,11^. The polar and charged atoms of the solute can hydrogen bond with water. Around apolar non-hydrogen bonding regions, in contrast, interfacial water molecules preferentially orient towards bulk water to maintain enthalpically favorable hydrogen bonds^12^. This counterbalances the entropic penalty for locally increasing the order of water molecules in the solvation shell, visible as spatial variations in density, enthalpy, and entropy. Another example of enthalpy-entropy compensation is the presence of water in hydrophobic cavities. Although the water trapped inside them is largely isolated from the bulk hydrogen-bonding network and its exchanges with bulk are slowed down^13^, its removal from the cavity would be unfavorable^14–16^.

In the literature, water at the interface with hydrophobic solutes and in hydrophobic pockets is termed *high-energy, hot*, or *unhappy* ^17^ and its arrangements are coined *clathrates* or *icebergs*^18–21^. Analogously, water in bulk is *low-energy, cold*, or *happy*. These concepts have motivated the identification and classification of water sites, recognizing water as an important determinant of thermodynamic stability. The energetics of displacing water from the binding pocket upon ligand binding are correlated with (a) the binding free energy^22–24^ or (b) transition state barriers and their resulting kinetic rates^25^. Several computational tools can help map local hydration thermodynamics in space^26–28^.

Molecular dynamics (MD) is a powerful computational technique that enables simulation of atomic movement over time, finding widespread application in diverse fields, among them biophysics. In ligand binding problems, simulations give critical atomistic insights into solvent rearrangement around both ligand and target. In the prototypical trypsin-benzamidine case, for example, multiple water configurations were observed while the ligand seemingly remained in its crystallographic binding pose. These different hydration states not only yielded different free energies, but also significantly influenced the binding/unbinding mechanism and the corresponding kinetic rates^29,30^.

Statistically significant measurements of thermodynamical properties require MD simulations to extensively sample the phase space. However, barriers during ligand unbinding and water exchange between bulk and solvation shell often create kinetic bottlenecks that reduce the frequency of events observed within the timescale accessible to standard unbiased simulations, limiting their accuracy. Monte Carlo equilibration of water has emerged as a promising approach to tackle the water sampling challenge^31–33^. Alternatively, collective variables-based enhanced sampling methods such as Metadynamics or On-the-fly Probability Enhanced Sampling (OPES) overcome the sampling problem by learning a bias potential that helps escape energetic minima^34–36^. These methods rely on the definition of collective variables (CVs), functions of atomic coordinates, whose sampling is to be accelerated. An effective choice of CVs that capture the slow degrees of freedom in a system is paramount for the success of enhanced sampling simulations^37^. Nonetheless, such an optimal choice is usually far from trivial.

Common CVs in ligand binding often are purely geometric, such as distances, contact maps, and angles. These choices are straightforward to implement but limited, as they accelerate only the solute, thereby neglecting the crucial role of the solvent. Effectively, using these water-blind CVs implicitly assumes a very fast adaptation of the surrounding water to the solute motion. The validity of this premise is challenged by the role of slowly diffusing water molecules. While the notion of accelerating the sampling of such critical water molecules is compelling, their precise identification persists as a considerable challenge that deters from realizing the full potential of dedicated hydration CVs.

Current strategies follow chemical intuition and choose to accelerate the water coordination around i) real atoms, such as geometrically distant polar atoms^38–41^ and ions^42^, ii) notable positions such as within hydrophobic pockets or along the binding axis^43^, or iii) in positions where water exhibits a long lifetime^29^. The explicit inclusion of hydration CVs successfully improved binding free energy predictions in a number of examples^43–48^.

In this work, we take inspiration from a solvent structure-based tool that quantifies as a fingerprint value the deviations from the uniform radial distribution function of water oxygen atoms^49^. The fingerprint scale is relative, and its numerical range may vary between systems. Therefore, we define its limits as *bulk* and *anti-bulk* according to the measured absolute minimum and maximum values for each system, respectively. We let the fingerprint guide us to pick two distinct sets of points of interest (POIs) that either give prominence to bulk-like or anti-bulk-like solvation. We then define hydration CVs as the water coordination around the POIs within each set. These two sets are separately biased in enhanced sampling simulations with the aim of comparing and assessing which strategy is the most effective.

We test our approach on a set of host-guest systems from the SAMPL5 and SAMPL6 challenges that are typically used as surrogates for protein-ligand systems, and where properly sampling the water motion around the ligands and the host hydrophobic cavity is a known challenge^50^. For reference, we use our recent simulations with OneOPES, a powerful method that combines OPES, replica exchange, multi-CV enhanced sampling and a funnel-shaped restraint^39^, that have achieved excellent agreement with experiments.

Here, we present a simpler data-driven strategy that is exclusively water-focused and bypasses heuristics in selecting hydration CVs. We validate the results corresponding to the two different selection criteria against the OneOPES references. The comparison with well-converged computational results eliminates force field-related limitations that could confound the attribution of discrepancies to insufficient sampling.

Our investigation is structured into two parts. First, we select water POIs at both extremes of the fingerprint scale and compute binding free energies in single replica OPES simulations where the water coordination around the POIs are the only CVs used. Remarkably, we find that such CVs alone are sufficient to converge the free energies, with the choice of anti-bulk-like POIs clearly emerging as the more effective. Typically, anti-bulk POIs lie deep inside the hydrophobic cavity of the host or correspond to apolar and uncharged, or buried atoms of the guest. Second, we compare results with those obtained from a standard choice of geometric CVs. Strikingly, we observe that biasing fingerprint-guided hydration CVs alone exceeds the performance of such geometric, water-blind CVs, achieving a faster and more robust convergence.

Altogether, this work illustrates the power of accelerating water using rationally and automatically picked hydration CVs. Although the choice to bias exclusively water coordination may appear counterintuitive at first sight, it demonstrates the added and distinctive value of accounting for hydration instead of entirely relying on water-blind descriptors. While larger, more flexible and more complex systems may benefit from the development of a hybrid hydration-geometric CVs approach, the fingerprint-driven identification of specific atoms and points in space where to bias water coordination still provides a critical stepping stone. This work opens the door to the systematic incorporation of hydration CVs in enhanced sampling, extending their routine application beyond ligand binding to all processes in solution.

## II. METHODS

All molecular dynamics simulations were carried out in GROMACS 2023^51^ patched with PLUMED version 2.9.1^52,53^. Supramolecular host TEMOA (tetra-*endo*-methyl-octa-acid) and guest molecules were parametrized as described in reference^39^, using the option that gave the best agreement with experiments. We used the GAFF2 force field^54^, the dielectric electrostatic model and the TIP3P water model^55^. Further details on the protocol and its implementation are described in^39,53^. The equations of motion were propagated with a leapfrog integrator with a timestep of 2 fs. All bonds were constrained with the LINCS algorithm^56^. Short-range van der Waals and Coulomb non-bonded interactions were cut off at a distance of 1 nm. The particle-mesh Ewald method^57,58^ was used for long-range electrostatics. All production runs were performed in the NVT ensemble. The temperature was kept at 298 K with a velocity-rescaling thermostat^59^.

### A. Calculation of the Fingerprint

To properly capture the interaction of solvent with either guests or host, it is important to evaluate each of them in solution separately. A single solute molecule — either the host or the guest — was at first solvated alone in a cubic box. The length of the box vectors was defined as the maximum distance between atoms in the solute plus 6 nm. This ensured that the fingerprint was not affected by periodic boundary conditions artifacts for radial distances of up to 2 nm. Counterions were added to neutralize the solute’s net charge. Following energy minimization using steepest descent, the system was equilibrated in the NVT ensemble for 1 ns before running an unbiased production run of 100 ns for the guests and 200 ns for the host system in solution.

The fingerprint *S* of point *i* is calculated using the equation introduced by Pérez-Conesa et al.^49^

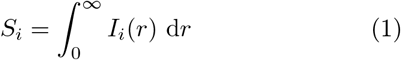

with the integrand being

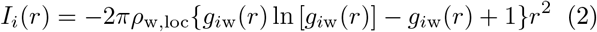

The local number density of water oxygen atoms *ρ*_w,loc_ and the radial distribution function of water oxygen atoms *g*_*i*w_(*r*) were extracted from the simulations. The radial distribution function at POI *i* was calculated up to a radius of 2 nm with bins of 0.001 nm width and a time resolution of 2 ps. GROMACS `gmx rdf` and PLUMED `COORDINATION` functions were used for guest (real) atoms and host virtual atoms, respectively. From the radial distribution function, *ρ*_w,loc_ was calculated as the average number density of the last 500 bins in radius range between 1.5–2.0 nm, as water is assumed to show bulk behavior there. The fingerprint was integrated from the origin to *r*_MAX_ = 2.0 nm using the trapezoid rule. The scripts that we used to generate the local fingerprint from simulations of small molecules in solution and select atoms around which to build hydration CVs are provided at https://github.com/valeriorizzi/WaterFP. A flowchart of the automatic selection process is shown in Figures S1 and S2.

### B. Enhanced Sampling Simulations

Here, we run enhanced sampling simulations using a recent evolution of Metadynamics^60,61^, the On-the-fly Probability Enhanced Sampling Method (OPES)^36^ that iteratively builds the bias from an on-the-fly estimation of the probability distribution. We use its Explore variant that is particularly suited when biasing more than two CVs^35^. For simplicity, in all simulations we use a BARRIER of 50 kJ/mol and a PACE of 10000 steps (20 ps). SIGMA values are automatically determined from the initial short unbiased portion of the trajectories.

We test three different simulation scenarios using different CV sets referred to as (a) *bulk fingerprint (fp)*, (b) *anti-bulk fp*, (c) *geometric*. In sets (a) and (b), the bias is applied exclusively to water in proximity of atoms selected based on their fingerprint, and in (c) we bias water-blind common geometric descriptors in host-guest systems. For each CV set, three independent simulations each of 500 ns are run for every pair of host TEMOA and guest of the SAMPL5^62^ and SAMPL6^63^ challenge. The input run files were taken from the GitHub repository by Febrer et al.^39^, while the PLUMED input files were adapted for the fingerprint and geometrical CV sets. These files are available at https://github.com/valeriorizzi/WaterFP. Common to all simulations is a funnel-shaped static potential^64,65^ that helps to reduce the configuration space accessible to the ligand that in the unbound state is restrained to a cylindrical volume of radius 2 Å.

In the two simulation sets (a) and (b), we employ three hydration CVs to build the external bias potential with OPES Explore. A hydration CV for POI *i* corresponds to the coordination number CN_*i*_ of *i* by water oxygen atoms defined by the switching function

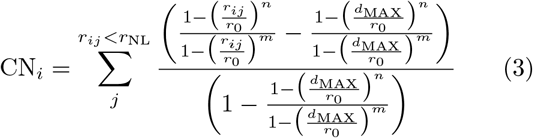

where *r*_*ij*_ is the distance between the POI *i* and the water oxygen atom *j*, summed over all the atoms within a sphere of radius *r*_NL_ = 16 Å. The list of oxygen atoms is updated every 20 steps. We set *r*_0_ = 3.5 Å and *d*_MAX_ = 12 Å, beyond which the function is zero. Following the considerations in Ref.^43^, {*m, n* } are {6, 10} and {2, 6}for real and virtual atoms, respectively. The bias potential mostly affects the water molecules in the vicinity of *i* and its effect softly decays toward distant ones. Of the three hydration CVs, one is always on the host virtual atom (v1) which has the lowest fingerprint value among the points tested, while the other two CVs are on either the two highest- or lowest-scoring fingerprint atoms of the ligand (see also Figure S3). If two top-ranking atoms were directly neighboring, the choice was adapted to pick two more distant atoms. We call these simulation sets *bulk fp* and *anti-bulk fp*, respectively.

In the last simulation set (c), we biased three water-blind geometric CVs: (i) the distance between host and guest *z*, (ii) the cosine of the angle between a ligand axis and the funnel axis *COS*, and (iii) the radial position of the ligand with respect to the funnel axis *r*_funnel_. We call this simulation set *geometric*.

These three sets of simulations are compared to those reported in Ref.^39^, which were computed with OneOPES^66^, a multi-replica, multi-CVs variant of OPES Explore^35^. These references were run as triplicates for 150 ns, using a funnel restraint and biasing *z* and *COS* as main CVs as well as a set of auxiliary hydration CVs picked heuristically.

### C. Uncertainty Quantification

We quantified the uncertainty of the predicted binding free energy at different simulation time lengths as follows. Three independent simulations were truncated each to the specified length while the first 10 ns were considered equilibration and discarded. The binding free energy was calculated via reweighting along the funnel coordinate *z* using the deposited bias. The average and the standard deviation of the three simulations were used to estimate the binding free energy and the associated uncertainty, respectively.

In addition, we calculated the coefficient of determination (R^2^) and the root mean square error (RMSE) to evaluate the agreement between the results obtained here with the different CVs sets and the reference OneOPES simulations^39^. The confidence limits for each statistical estimator were calculated via bootstrap by resampling the estimators 10^5^ times using a script published by Rizzi *et al*.^67^.

## III. RESULTS

We begin by characterizing the water in our system with the fingerprint tool. While the strategy does not require knowledge on the binding pose and any point in space could be picked, we focus our analysis of water behavior on the most relevant points for the transition between the bound and unbound state, thus reducing the computational cost. Turning to the host first, water molecules inside the host cavity are displaced upon binding. Therefore, we calculate the fingerprint value for virtual atoms along the central molecular axis of the host (*r*_funnel_ = 0 nm, the *C*_4_ symmetry axis). This allows capturing approximately the cavity volume with spherical descriptors. For the guests, we calculate the fingerprint for all their heavy atoms.

Figure 1 shows the radial distribution function of the water oxygen atoms *g*_*i*w_(*r*), the corresponding fingerprint value *S*_*i*_, and its integrand *I*_*i*_(*r*) for the virtual atoms v1 and v2 of host TEMOA and atoms C8 and O1 of SAMPL5 guest G2. These atoms have the absolute maximum and minimum fingerprint values within the respective molecules. In TEMOA, the host virtual atom v1 is located in the center of the hydrophobic cavity. Here, *g*_v1w_(*r*) peaks at *r* ≈ 0.15 nm indicating the presence of trapped water molecules. As expected, *g*_v1w_(*r*) vanishes in the region occupied by the host atoms. This region gives the largest contribution to the fingerprint value. At larger distances, *g*_v1w_(*r*) fluctuates around unity (dashed line).

**FIG. 1:**
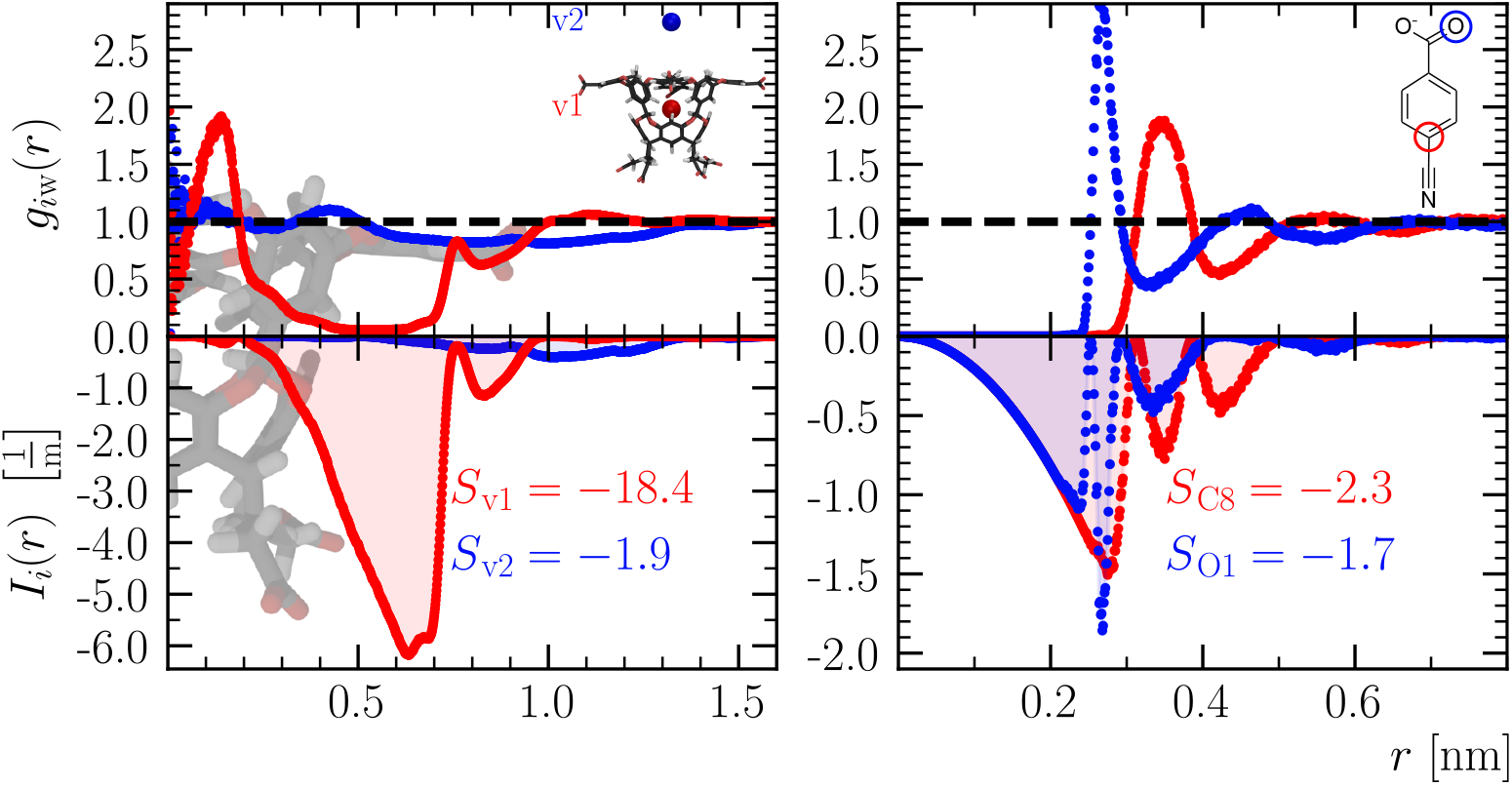
Radial distribution function *g*_*i*w_(*r*), integrand *I*_*i*_(*r*) and fingerprint *S*_*i*_ for (left) selected virtual atoms of the TEMOA host and (right) selected real atoms of the S5-G2 guest.

Picking a POI at a larger distance from the cavity, shown here for v2, the perturbative effect of the solute decreases, and thus *g*_v2w_(*r*) does not deviate much from unity. In agreement with this, *S*_v2_ decreases much less than *S*_v1_.

For guest atoms such as in molecule S5-G2, due to the repulsion between a real atom and the surrounding water molecules, *g*_*i*w_(*r*) is zero up to the first solvation shell where it reaches its maximum value. For the polar oxygen atom O1, *g*_*i*w_(*r*) peaks earlier than for the apolar carbon atom C8 in S5-G2. Deviations of *g*_*i*w_(*r*) from unity contribute to the fingerprint value with the cube of the radius. Consequently, the fingerprint is sensitive to the solvent exclusion radius and increases with the radial distance of the perturbation.

We repeat the same analysis for each non-hydrogen atom in the guests, leveraging the advantage of the fingerprint computation being easily automated. The results are reported in Figure 2. Polar atoms tend to cluster at one end of the fingerprint scale, whereas apolar atoms accumulate at the opposite end. Notably, the same chemical element can have varying fingerprint values depending on its molecular environment. In particular, buried atoms tend to have very large fingerprint values due to their significant exclusion volume.

**FIG. 2:**
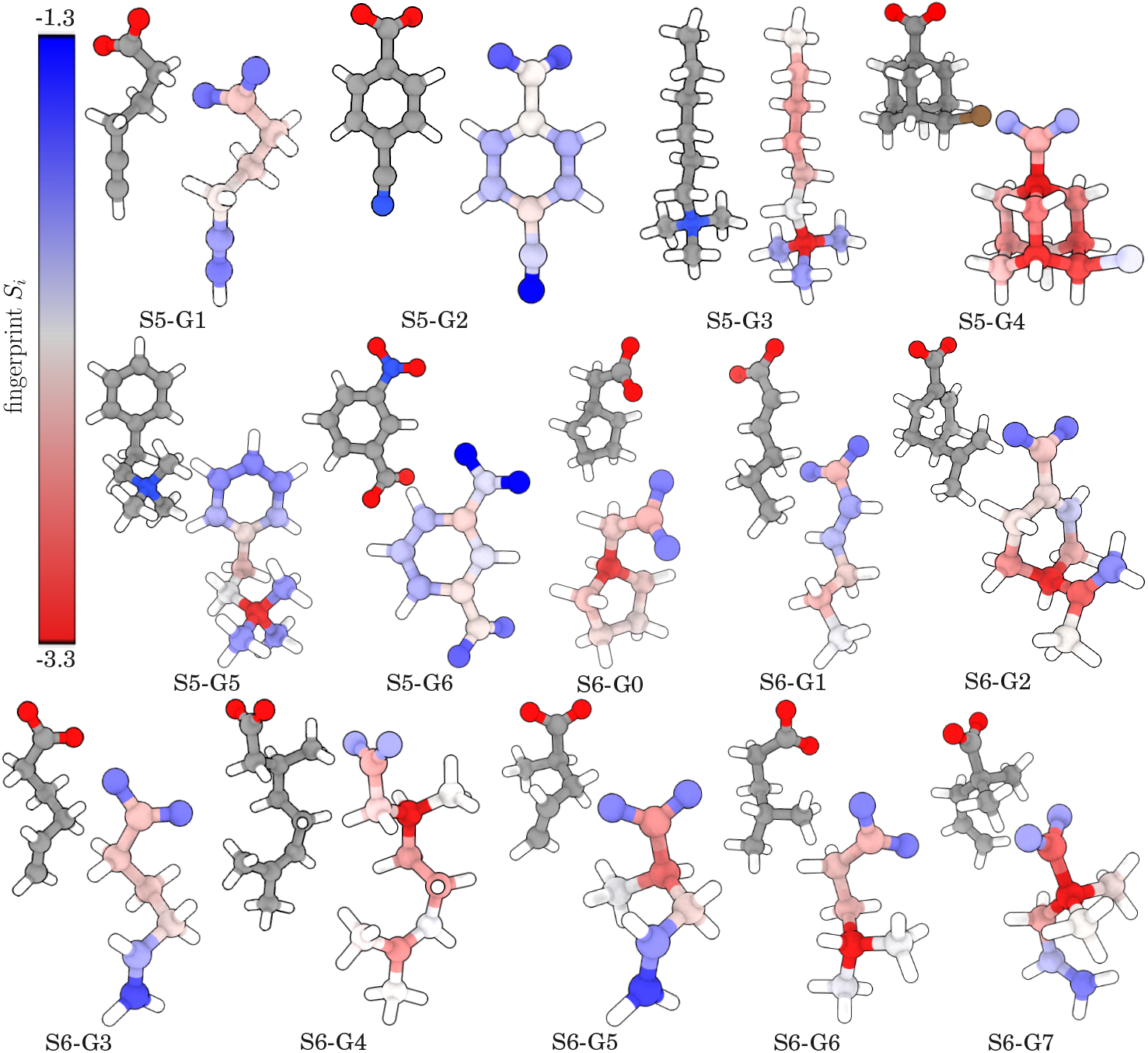
Global comparison of fingerprint values of SAMPL5 and SAMPL6 guest molecules. Each guest molecule is shown in two representations. On the left, its atoms are colored according to their element type (oxygen, nitrogen, carbon, and hydrogen atoms are colored in red, blue, gray and white, respectively). On the right, each atom *i* of a guest molecules is colored according to its fingerprint value *S*_*i*_, except for hydrogen ones, which are only shown for clarity.

Yet, this correlation between solvent excluded volume and fingerprint value is not trivially linear or even monotonic. To better quantify the effect of steric shielding and differentiate it from the local solvent effect captured by the fingerprint, we calculate the Solvent Accessible Surface Area (SASA) for all heavy atom of the ligands (see Figure S26) and we show its correlation with fingerprint values in Figure S27. We single out ligands S6-G1 and S6-G6 as two opposing examples: while for S6-G6 the correlation SASA-fingerprint is strong, ligand S6-G1 shows instead a poor corelation, especially for atoms C1 and C2 (see Figure S28). These latter atoms have a low SASA value, yet their fingerprint values are closer to those of more bulk-like atoms. Since the host TEMOA usually binds to apolar moieties and a charged or polar head sticks out of the binding pocket, the anti-bulk and bulk fingerprint atoms can be mapped to these regions, respectively.

To test which POIs identified by the fingerprint are best suited for hydration CVs to be biased, we directly compare the choice of bulk and anti-bulk fingerprint atoms, as shown in Figure S3. The quality of each choice is evaluated by comparing the resulting free energy estimates with reference and experimental values. The results are shown in Figures 3 and S23 and summarized in Table S1 and Table S2, respectively. An outlier system, S5-G4, was removed from the dataset because of convergence issues that we discuss below.

**FIG. 3:**
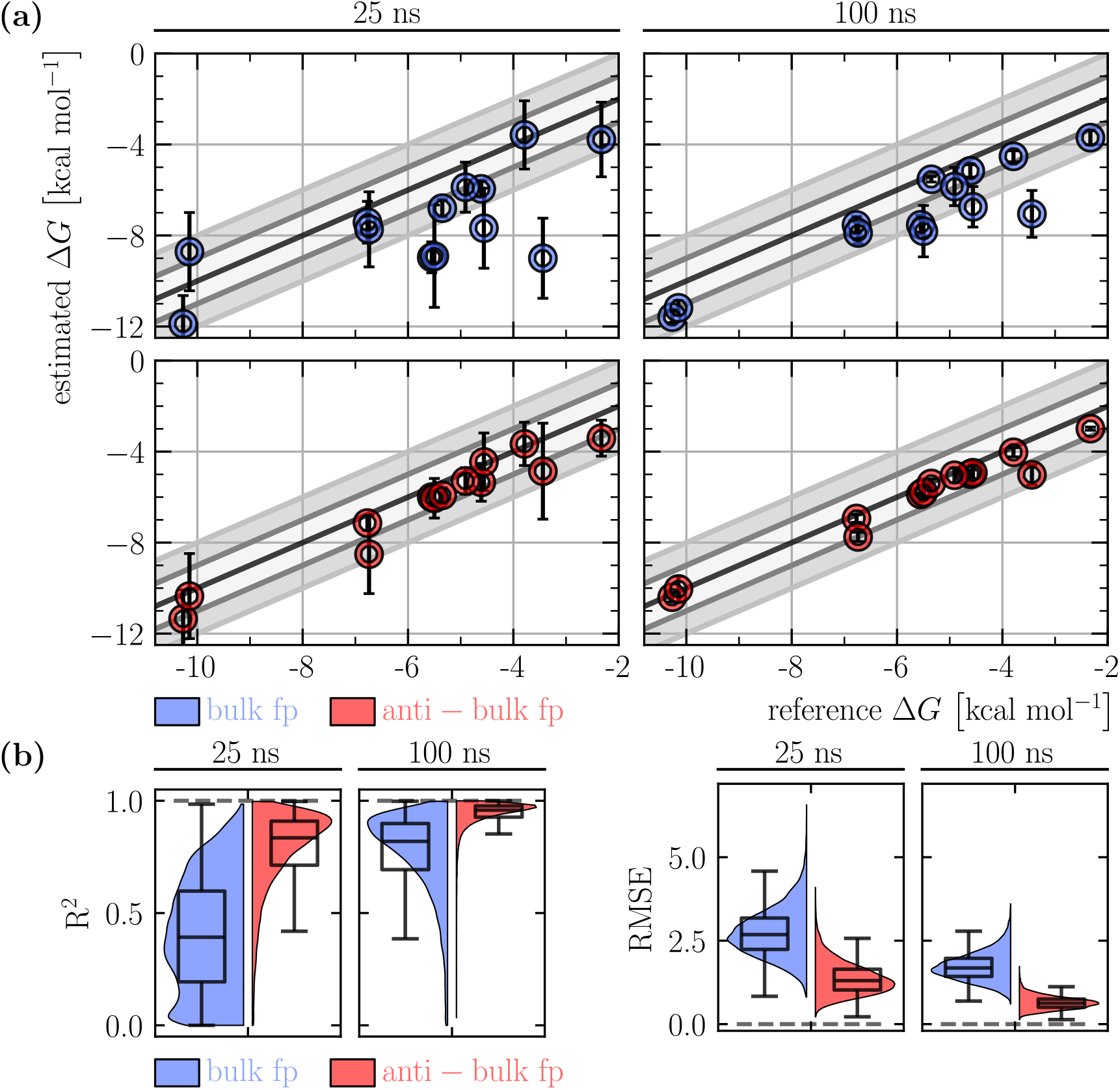
(a) Comparison of free energy estimates at 25 and 100 ns for bulk (blue) and anti-bulk (red) fingerprint CVs to the reference free energies. Error bars show one standard deviation. (b) Statistical estimators of the agreement in (a) and their bootstrapped distributions.

For shorter simulation times, choosing bulk fingerprint atoms in the guest resulted in poorer convergence and, consequently, worse agreement with the reference. The difference between bulk and anti-bulk fingerprint atoms gradually decreases with increasing simulation time, suggesting that the choice of hydration CVs influences the speed of convergence rather than the capability to converge to the reference binding free energy values themselves (see also Figures S4–S17).

The finding that fingerprint-guided hydration CVs alone are able to produce converged free energy results for a large number of host-guest systems prompted us to compare the top performing set of hydration CVs to the popular choice of geometric CVs that is entirely water-blind. As shown in Figure 4, the binding free energy estimates at a simulation length of both 25 ns and 100 ns reveal that the anti-bulk hydration CVs results are in better agreement with the reference than the geometric descriptors. In fact, obtaining equivalent results using the geometric CVs requires much longer simulations (≳ 200 ns, see Figure S20), highlighting water as a more crucial slow mode to be accelerated. The time evolution of the statistical estimators in Figure 4b mirrors the better performance of the hydration CVs (see also Figure S18 and Figure S19). Comparison to experimental values at short and long simulation lengths are shown in Figures S23, S24, and S25.

**FIG. 4:**
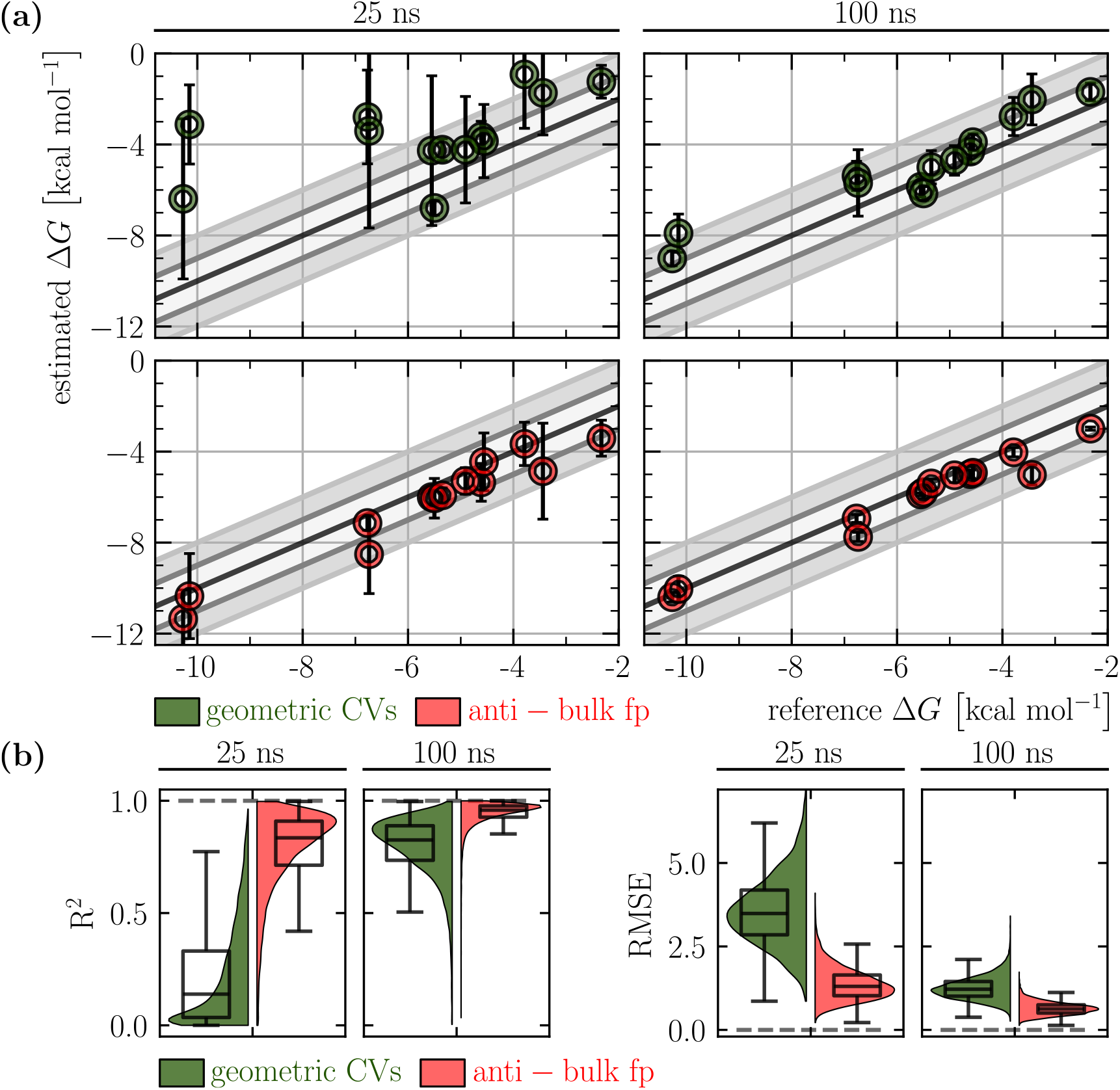
(a) Comparison of free energy estimates at 25 and 100 ns for geometric CVs (green) and anti-bulk fingerprint CVs (red) to the reference free energies. Error bars show one standard deviation. (b) Statistical estimators and their bootstrapped distributions.

To better dissect and illustrate the sampling differences between the three CVs sets, we analyze in more detail the binding free energy as a function of time for three representative systems. The results are reported in Figure 5, and further details for the other host-guest combinations are collected in Figures S21 and S22. In systems like S5-G1 and S5-G2, the anti-bulk hydration CVs and the geometric CVs yield equivalent results at 100 ns, while bulk hydration CVs produce a shifted binding free energy estimate. Systems such as S6-G0 seem less sensitive to the choice of hydration CVs, yet geometric CVs take longer to converge.

**FIG. 5:**
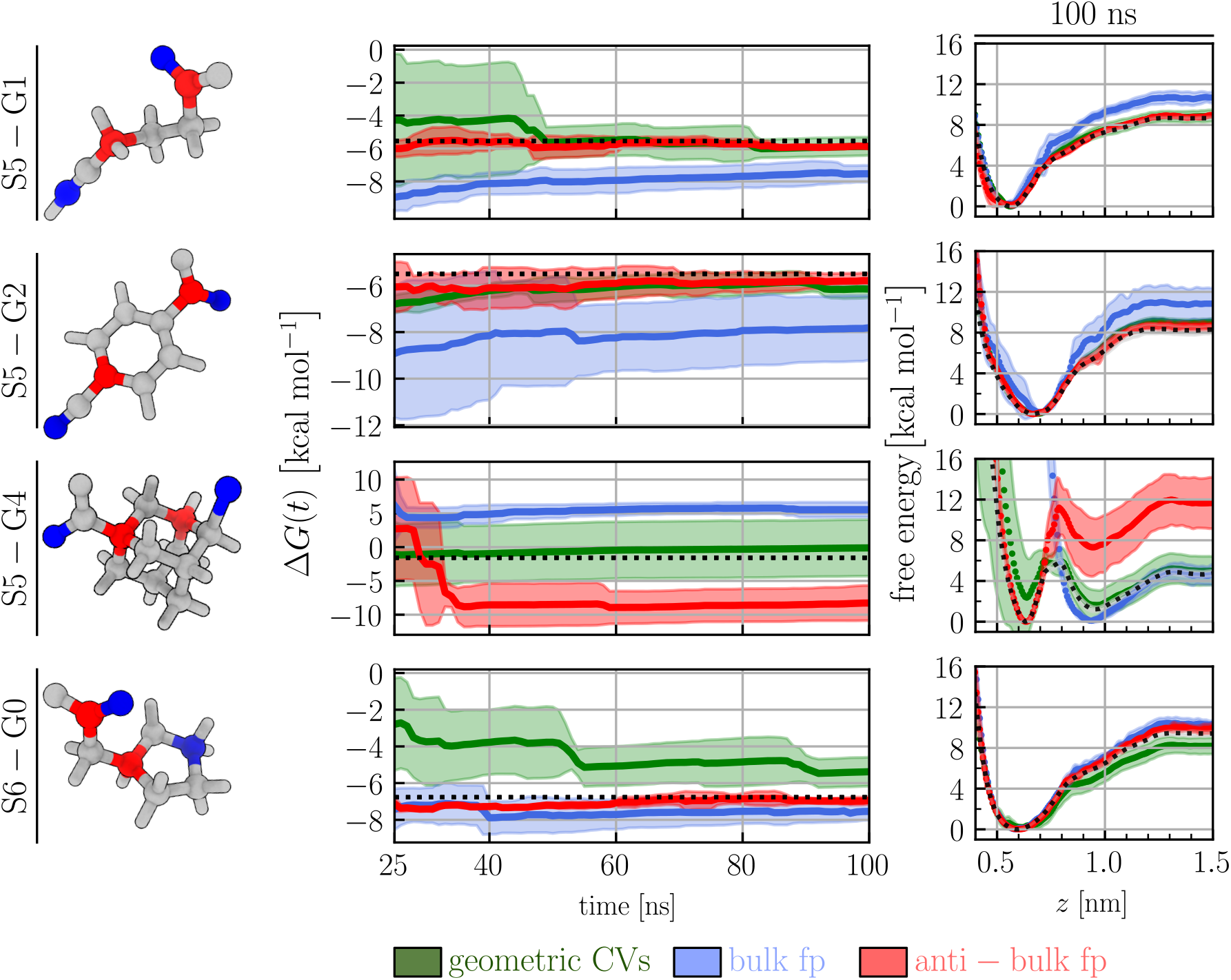
In the first column, the atomistic structure of the S5-G1, S5-G2, and S6-G0 guests. The colors indicate the atoms biased in the bulk (blue) and anti-bulk (red) CVs sets. In the second and third columns, the binding free energy estimates as a function of simulation length and the free energy profiles at 100 ns of sampling, respectively, for all three CVs sets. The simulations averages are shown as thick lines while the standard deviation as correspondingly colored areas. The dotted black lines show the reference values. The reference standard deviations are within line width in the second column and are shown in light gray in the third one.

The S5-G4 system is instead the only example in our test set where biasing only the hydration CVs is no longer sufficient. Previous work^43,68^ highlighted that this system has two competing bound states, as well as two metastable intermediate states. Both the bound states and the intermediate states present a water-rich and a water-depleted state. The similarity of water behavior between bound and intermediate states limits the ability of hydration CVs to navigate the configurational landscape and effectively drive intermediate to bound transitions, leading to convergence issues (see also Figures S7 and S21). In contrast, geometric CVs, while still ineffective in driving transitions between water-rich and water-depleted states, show better performance because they are able to distinguish intermediate states from bound states. This is a case where a hybrid approach that combines geometric and hydration CVs should be preferred.

The high quality of the fingerprint-guided anti-bulk hydration CVs is reflected by the quick convergence of the OPES bias potential to a quasi-static regime, overall yielding binding free energy estimates with a small standard deviation between independent replicas. Choosing a suboptimal set of CVs that neglects critical slow degrees of freedom requires longer simulation times, until the relevant degrees of freedom are sufficiently sampled. The caveat here is that the required amount of sampling is not known in advance.

## IV. DISCUSSION

We have presented a rational and automated strategy of hydration CV selection that consists of assigning a fingerprint value to POIs and then biasing corresponding hydration CVs on the top-scoring POIs. This strategy is computationally cheap and can reliably identify where water is an important slow degree of freedom to be accelerated in enhanced sampling in the presented set of host-guest systems. The automatic identification reduces human effort and decreases the need for chemical intuition or system-dependent knowledge in hydration CV selection.

Perhaps surprisingly, biasing water molecules around POIs with an anti-bulk fingerprint as the only CVs — without explicitly biasing any standard geometric CV for host and guest such as distances, angles or contact maps — quickly leads to robust converged binding free energies in the systems studied here. The choice of these POIs is in stark contrast to polar atoms selected by previous heuristic criteria, which roughly correspond to the case of bulk fingerprint POIs that we have shown to perform comparatively worse. Notably, the atoms of the two sets are often directly adjacent, yet one of the two sets clearly outperforms the other. This, and the fact that the same chemical element can have very different fingerprint values, highlights the crucial dependence on the local environment that the fingerprint encodes. We validate the generalizability and robustness of the strategy against standard geometric CVs in a large number of host-guest systems and consistently measure performance improvements.

Applying our strategy to larger systems would be a natural avenue that, however, may require further tests and improvements. The spherical description of the hydration CVs, in particular, would hamper the application of the fingerprint to large solutes, where the exclusion volume and not the solvent structure dominates the fingerprint. Furthermore, we expect that in larger and more complex systems, biasing hydration CVs alone may not be sufficient. Nevertheless, we believe that our conclusion that ignoring water as a crucial slow degree of freedom to be accelerated comes at the expense of convergence speed stands, regardless of the size and the complexity of a system. While improvements in generalizing the strategy are needed, we believe that defining powerful hydration CVs is even more pertinent for larger systems, as the long time limit can easily be out of reach.

All in all, we advocate for the importance of including hydration CVs and offer a strategy to leverage their full potential. Our findings hopefully stimulate their use in processes occurring in solution and the fingerprint value could be a useful quantity reflecting the solvent structure for Quantitative Structure-Activity Relationship (QSAR) models.

## SUPPLEMENTARY MATERIAL

The supplementary material collects further details on the automatic atom selection, the atoms biased in the bulk and anti-bulk simulations, the binding free energy for all the systems as a function of different simulations times, the corresponding 1D free energy profiles against the funnel *z* CV, further statistical descriptors to evaluate the agreement between predicted and reference free energy values, SASA and ligand connectivity correlation analysis, the fingerprint for different water models, and template scripts for OPES simulations.

## ACKNOWLEDGMENTS

The authors acknowledge the Swiss National Super-computing Centre (CSCS) for supercomputer time allocation (project ID: s1274 and lp84). They also acknowledge the Swiss National Science Foundation and Bridge for financial support (project numbers: 200021_204795, CRSII5_216587, and 40B2-0_203628). M.S. acknowledges the funding from the German Academic Exchange Service (DAAD) (scholarship ID: 57646565). The authors are grateful to Pedro Febrer Martinez for useful discussions.

## AUTHOR DECLARATIONS

### Conflict of Interest

The authors have no conflicts to disclose.

### Author Contributions

**Simone Aureli**: Methodology (equal); Writing – review & editing (equal). **Francesco Luigi Gervasio**: Funding acquisition (equal); Project administration (equal); Resources (equal); Supervision (equal); Writing – review & editing (equal). **Tetiana Khakhula**: Conceptualization (equal); Data curation (equal); Formal analysis (equal); Investigation (equal); Methodology (equal); Software (equal); Validation (equal); Visualization (equal); Writing – original draft (equal); Writing – review & editing (equal). **Nicola Piasentin**: Methodology (equal); Writing – review & editing (equal). **Valerio Rizzi**: Conceptualization (equal); Methodology (equal); Writing – review & editing (equal). **Marc Schulze**: Conceptualization (equal); Data curation (equal); Formal analysis (equal); Investigation (equal); Methodology (equal); Software (equal); Validation (equal); Visualization (equal); Writing – original draft (equal); Writing – review & editing (equal).

## DATA AVAILABILITY

At https://github.com/valeriorizzi/WaterFP we provide the PLUMED input files to run all the simulations with the bulk, anti-bulk, and geometric descriptor CVs. We also provide the corresponding free energy profiles for all the systems and scripts to automatically generate the local fingerprint from a simulation of a small molecule in solution. For all the GROMACS inputs and for further details on the system parametrization, please check the original works^39,53^ from which they were taken. They are made available at the corresponding GitHub repository at https://github.com/Pefema/OneOpes_protocol. Further details and data are available from the corresponding author upon reasonable request.

## Notes

https://github.com/valeriorizzi/WaterFP

### Competing Interest Statement

The authors have declared no competing interest.

### Summary of Updates

General revision of the text and the figures

