## Supporting Information for "All You Need Is Water: Converging Ligand Binding Simulations with Hydration Collective Variables"

### Fingerprint-guided atom selection

In this section, we first report a schematic pseudocode to describe the procedure of atom selection based on the calculated fingerprint values. The full code is reported in the Github repository at https://github.com/valeriorizzi/WaterFP. We then show in Figure S3 the bulk and anti-bulk atoms we have selected. For the sake of clarity, atoms with the highest and lowest fingerprint scores were assigned as bulk (blue) and anti-bulk (red), respectively. For atoms belonging to the same functional group (e.g., oxygen in the carboxylic acid functional group of S5-G2), only one of the two degenerate atoms has been selected.

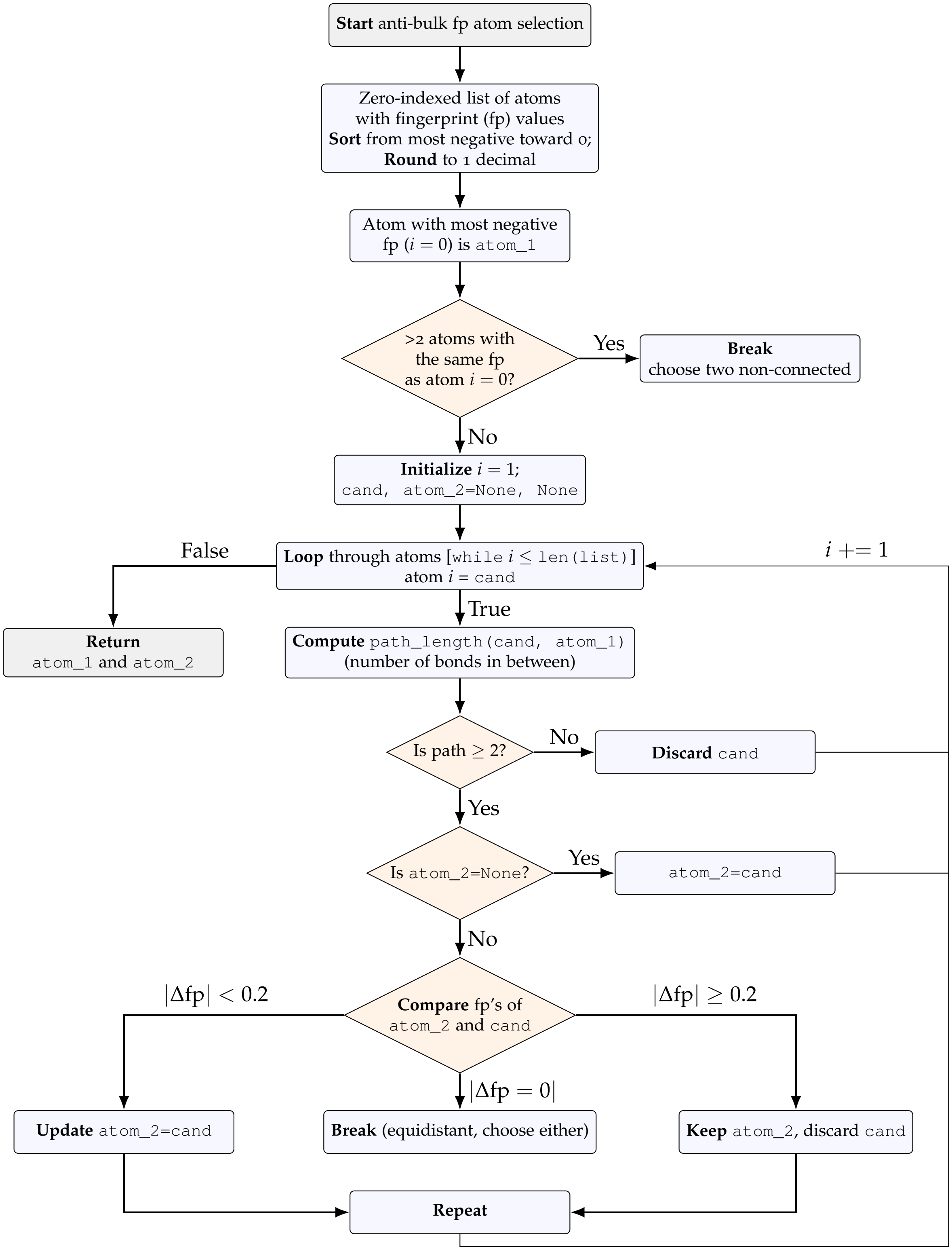


Figure S1: Flowchart showing fingerprint-based automatic selection of anti-bulk-like fp atoms in the guest molecules.

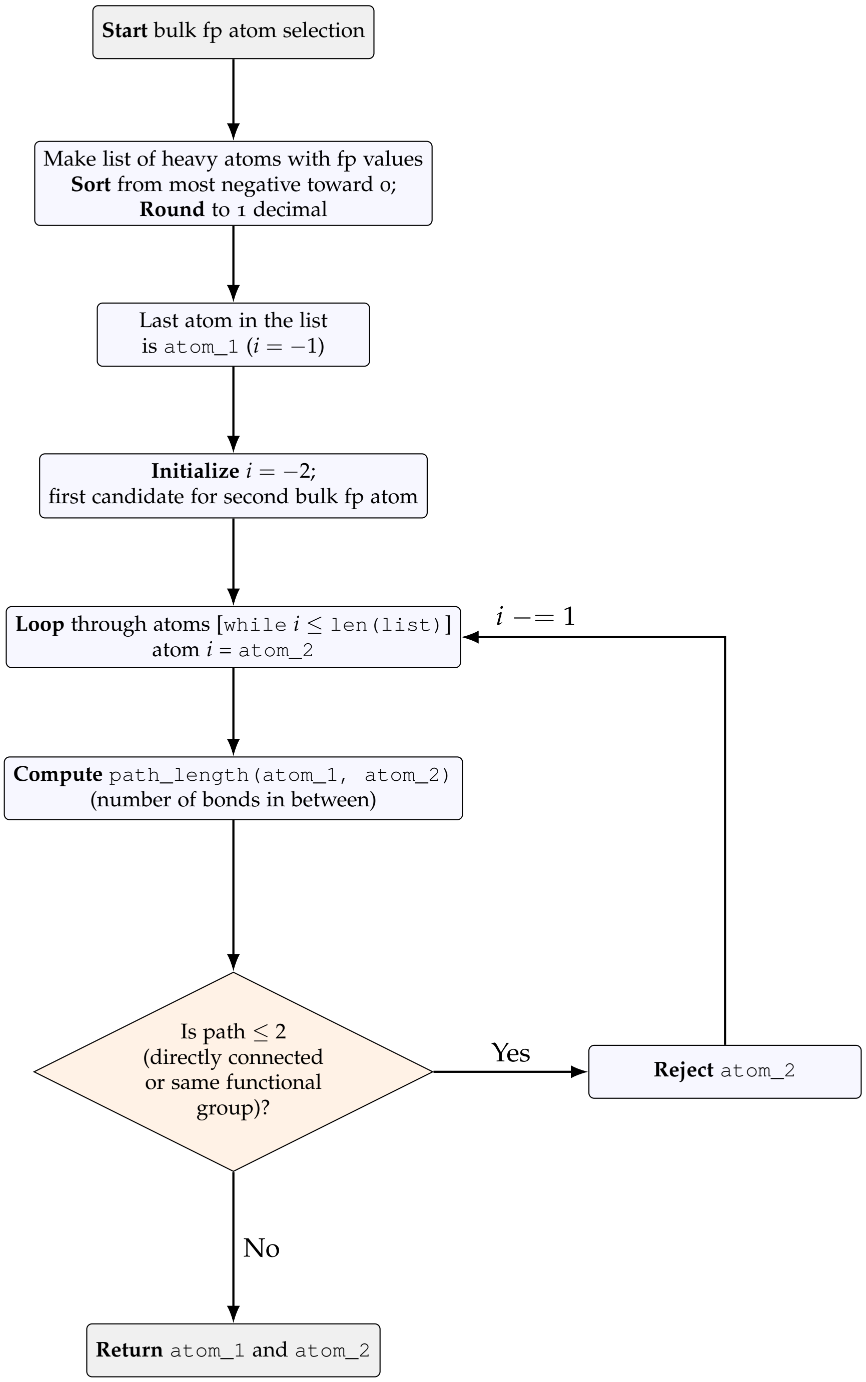


Figure S2: Flowchart showing Flowchart showing fingerprint-based automatic selection of bulk-like fp atoms in the guest molecules.

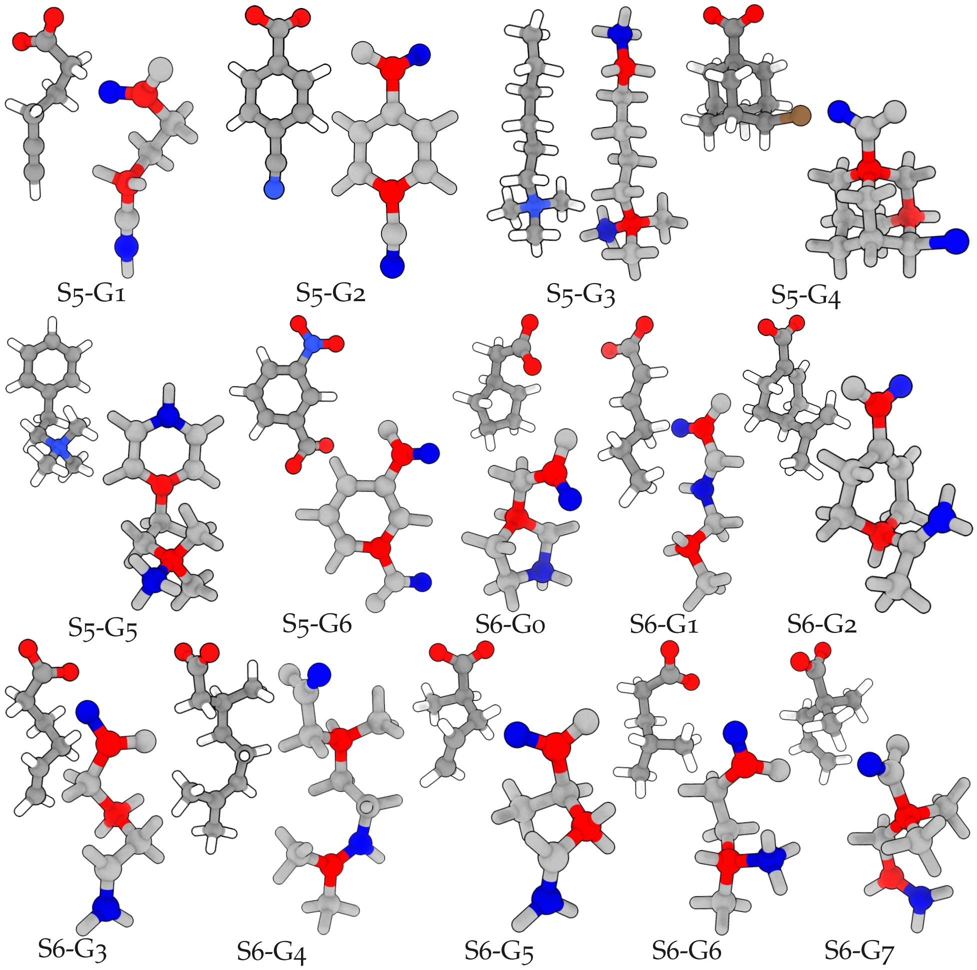


Figure S3: The two guest atoms selected to bias water in the case of bulk (blue) and anti-bulk (red) fingerprint CVs for all the guests simulated. The atoms of the insets are colored according to their chemical nature. Oxygen, nitrogen, carbon, and hydrogen atoms are colored in red, blue, gray and white, respectively.

### Free energy profiles over time for all the ligands and CVs

In this section, we report additional details of the OPES binding simulations carried out on the TEMOA host molecules and the ligands S5-G1 (Figure S4), S5-G2 (Figure S5), S5-G3 (Figure S6), S5-G4 (Figure S7), S5-G5 (Figure S8), S5-G6 (Figure S9), S6-G0 (Figure S10), S6-G1 (Figure S11), S6-G2 (Figure S12), S6-G3 (Figure S13), S6-G4 (Figure S14), S6-G5 (Figure S15), S6-G6 (Figure S16), S6-G7 (Figure S17). For each host/guest system, we display the time-evolution of the binding free-energy profiles with the three different CVs sets and the exploration of the $z$ CV along the funnel-shaped restraint.

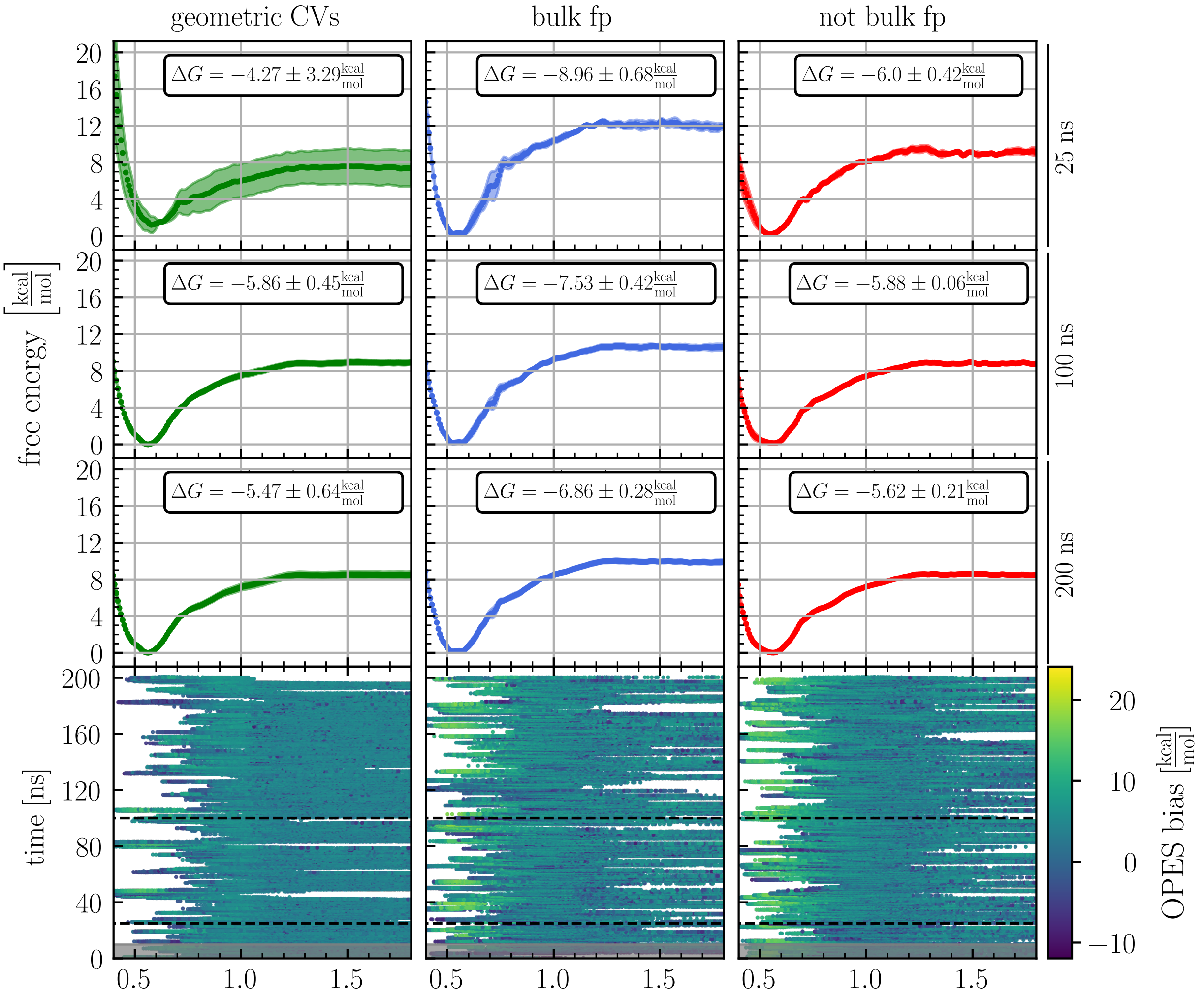


Figure S4: Free energy profiles as a function of $z$ and dynamics of $z$ as a function time for the host TEMOA and the guest S5-G1. The columns indicate the three different CVs sets. The panels in the first three rows show the averaged free energy profiles (dotted lines), their standard deviations (colored area), and the corresponding binding free energies $\Delta G$ after 25, 100, and 200 ns of sampling, respectively. The bottom row shows the time evolution of the $z$ funnel CV colored by the deposited bias. The first 10 ns shaded in gray are considered equilibration and discarded.

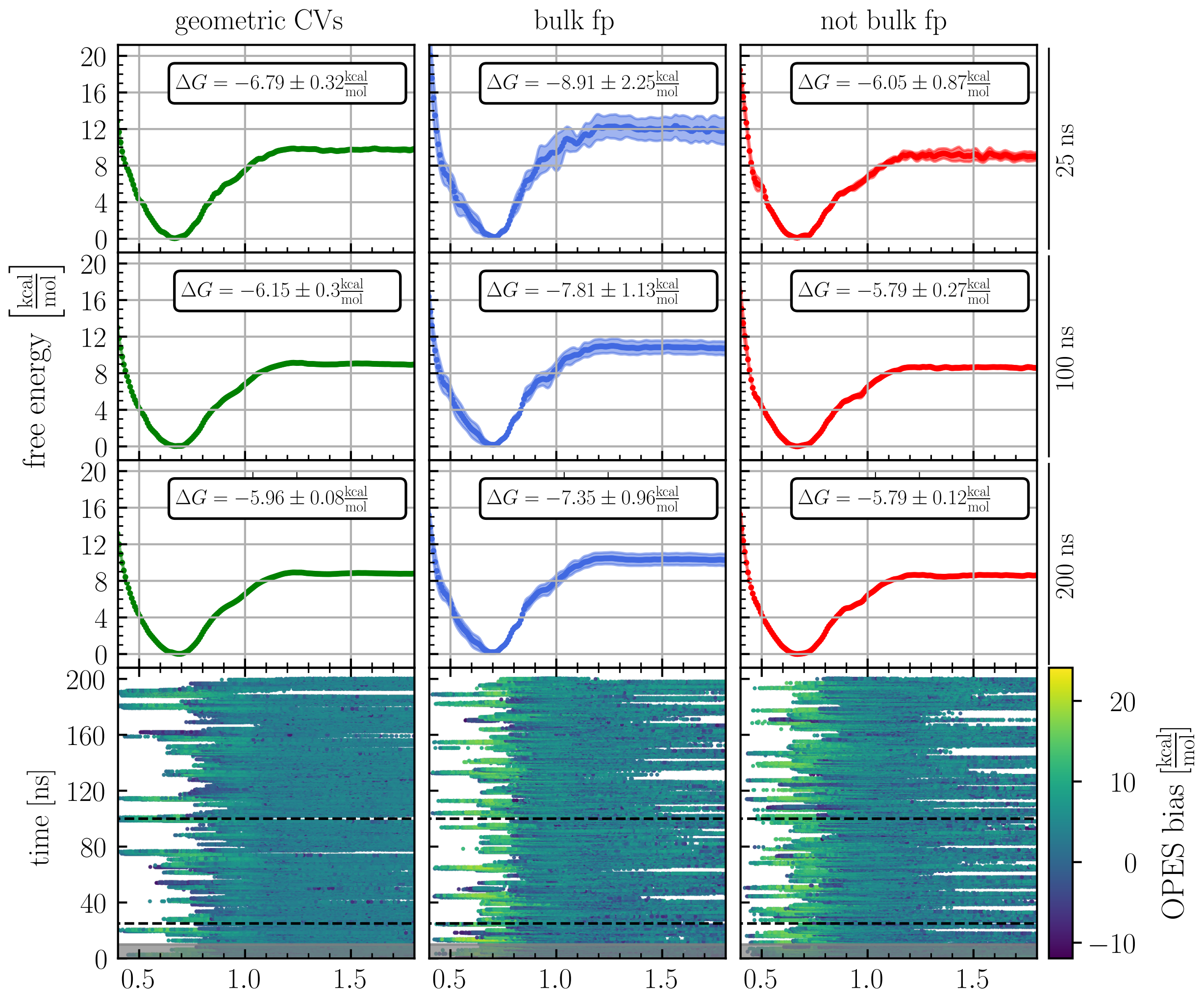


Figure S5: Free energy profiles as a function of $z$ and dynamics of $z$ as a function time for the host TEMOA and the guest S5-G2. The columns indicate the three different CVs sets. The panels in the first three rows show the averaged free energy profiles (dotted lines), their standard deviations (colored area), and the corresponding binding free energies $\Delta G$ after 25, 100, and 200 ns of sampling, respectively. The bottom row shows the time evolution of the $z$ funnel CV colored by the deposited bias. The first 10 ns shaded in gray are considered equilibration and discarded.

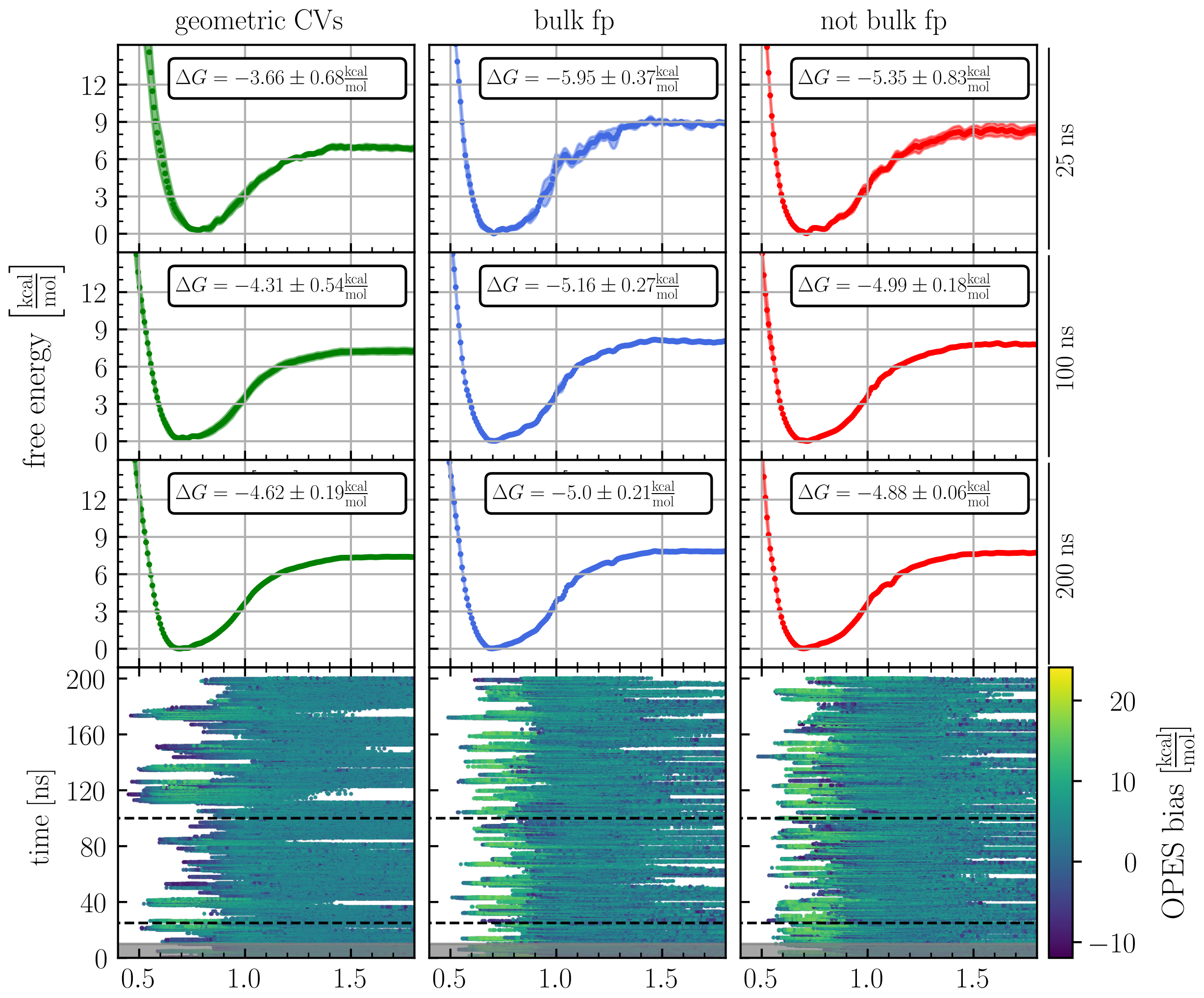


Figure S6: Free energy profiles as a function of $z$ and dynamics of $z$ as a function time for the host TEMOA and the guest S5-G3. The columns indicate the three different CVs sets. The panels in the first three rows show the averaged free energy profiles (dotted lines), their standard deviations (colored area), and the corresponding binding free energies $\Delta G$ after 25, 100, and 200 ns of sampling, respectively. The bottom row shows the time evolution of the $z$ funnel CV colored by the deposited bias. The first 10 ns shaded in gray are considered equilibration and discarded.

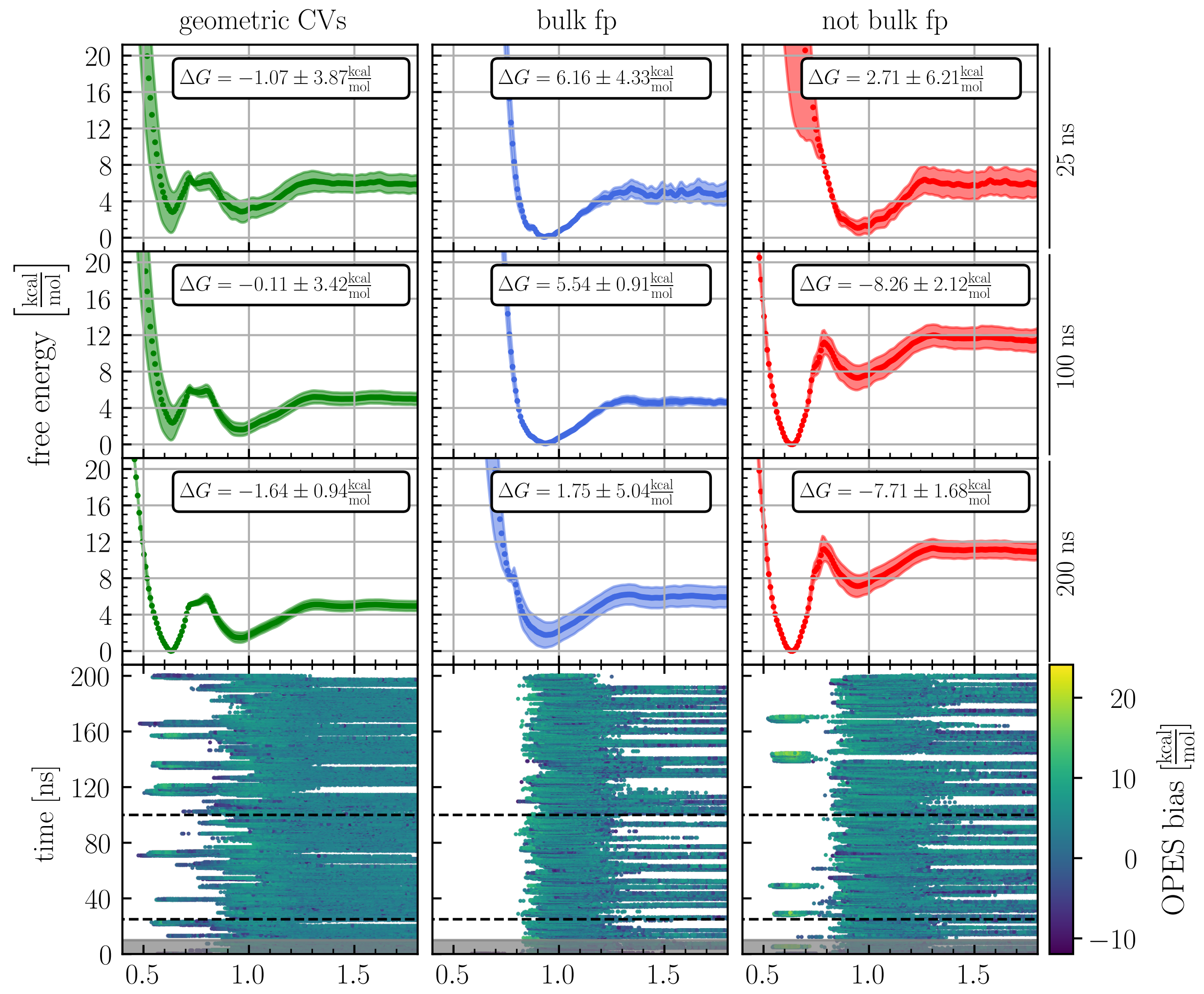


Figure S7: Free energy profiles as a function of $z$ and dynamics of $z$ as a function time for the host TEMOA and the guest S5-G4. The columns indicate the three different CVs sets. The panels in the first three rows show the averaged free energy profiles (dotted lines), their standard deviations (colored area), and the corresponding binding free energies $\Delta G$ after 25, 100, and 200 ns of sampling, respectively. The bottom row shows the time evolution of the $z$ funnel CV colored by the deposited bias. The first 10 ns shaded in gray are considered equilibration and discarded.

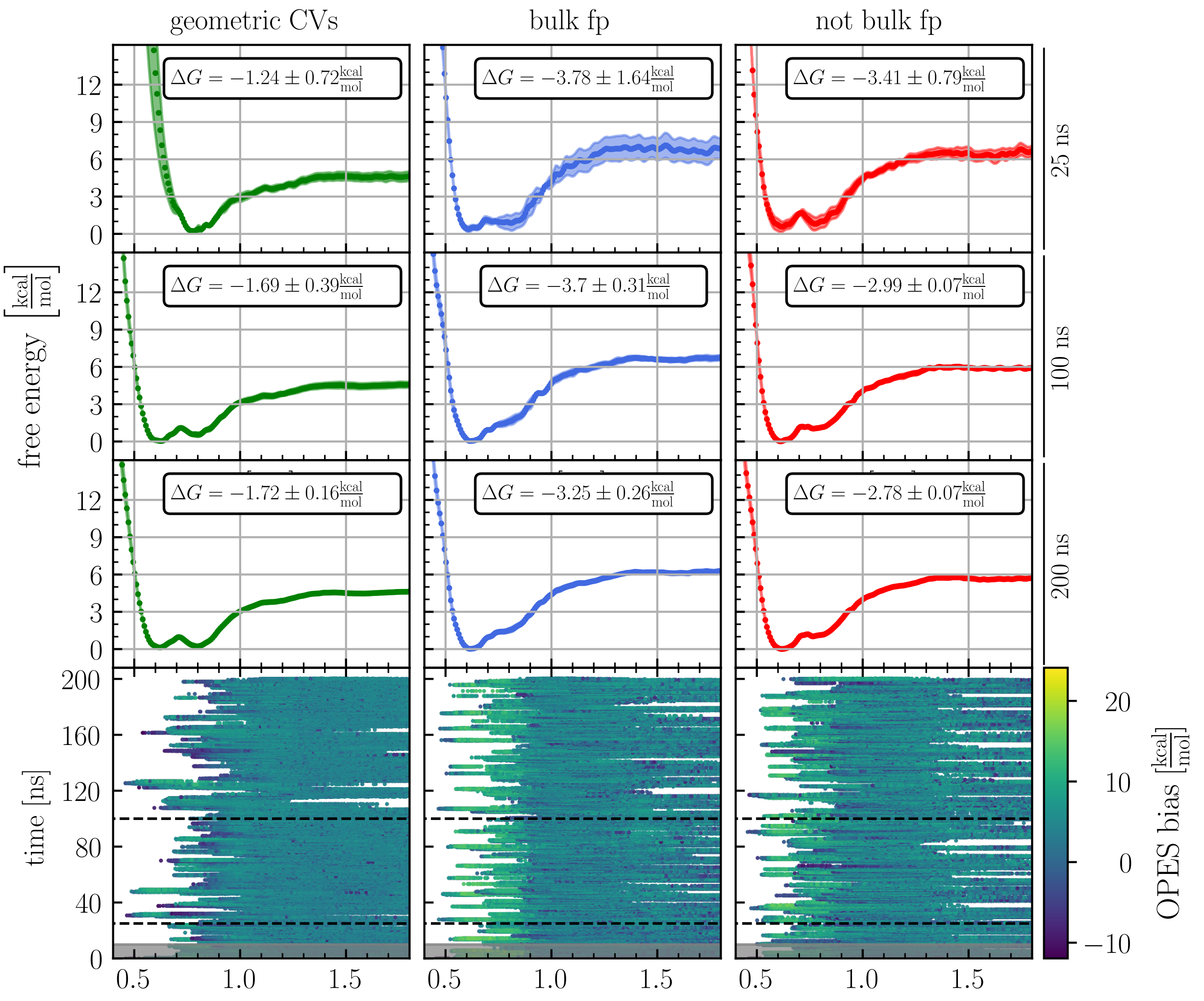


Figure S8: Free energy profiles as a function of $z$ and dynamics of $z$ as a function time for the host TEMOA and the guest S5-G5. The columns indicate the three different CVs sets. The panels in the first three rows show the averaged free energy profiles (dotted lines), their standard deviations (colored area), and the corresponding binding free energies $\Delta G$ after 25, 100, and 200 ns of sampling, respectively. The bottom row shows the time evolution of the $z$ funnel CV colored by the deposited bias. The first 10 ns shaded in gray are considered equilibration and discarded.

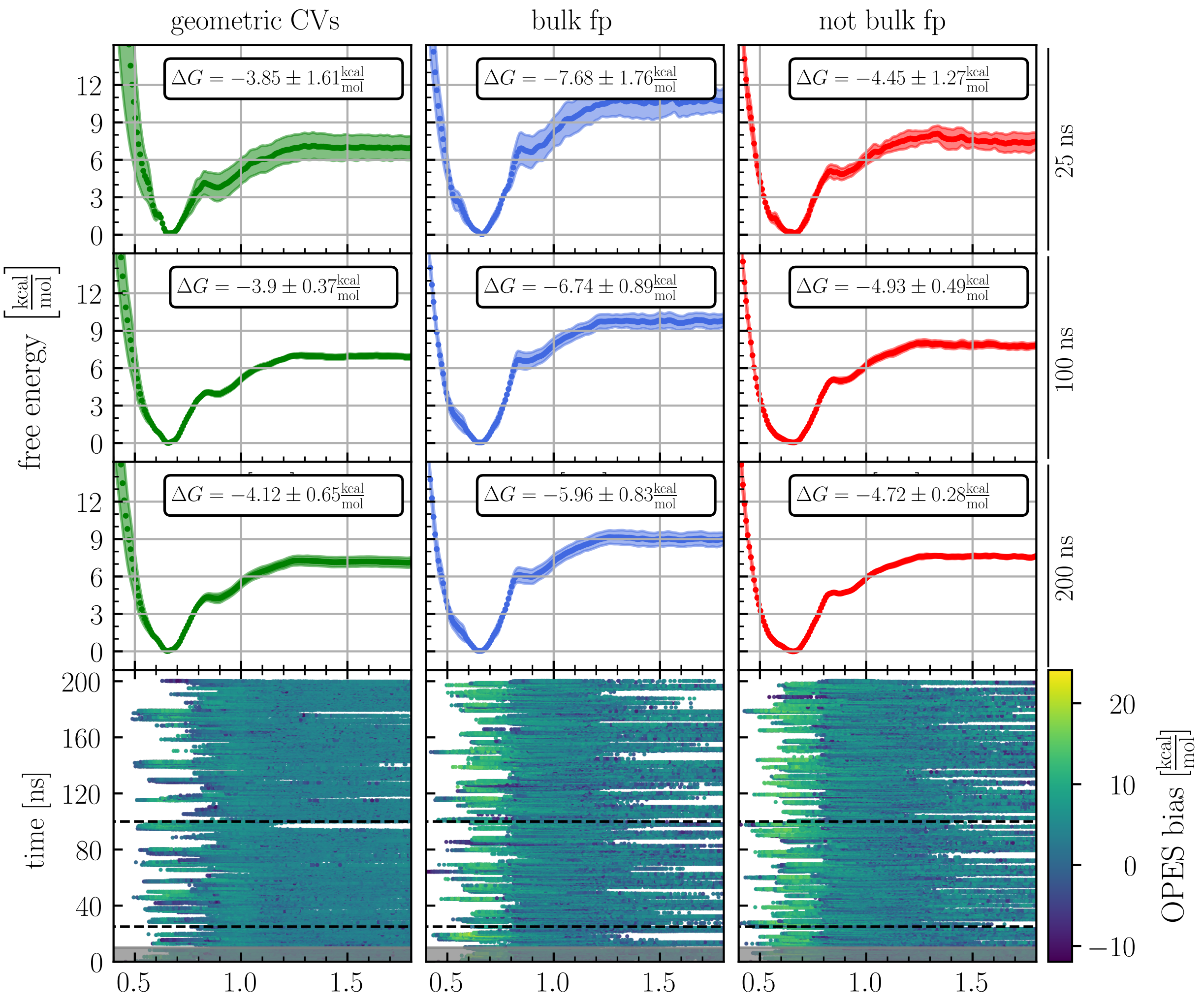


Figure S9: Free energy profiles as a function of $z$ and dynamics of $z$ as a function time for the host TEMOA and the guest S5-G6. The columns indicate the three different CVs sets. The panels in the first three rows show the averaged free energy profiles (dotted lines), their standard deviations (colored area), and the corresponding binding free energies $\Delta G$ after 25, 100, and 200 ns of sampling, respectively. The bottom row shows the time evolution of the $z$ funnel CV colored by the deposited bias. The first 10 ns shaded in gray are considered equilibration and discarded.

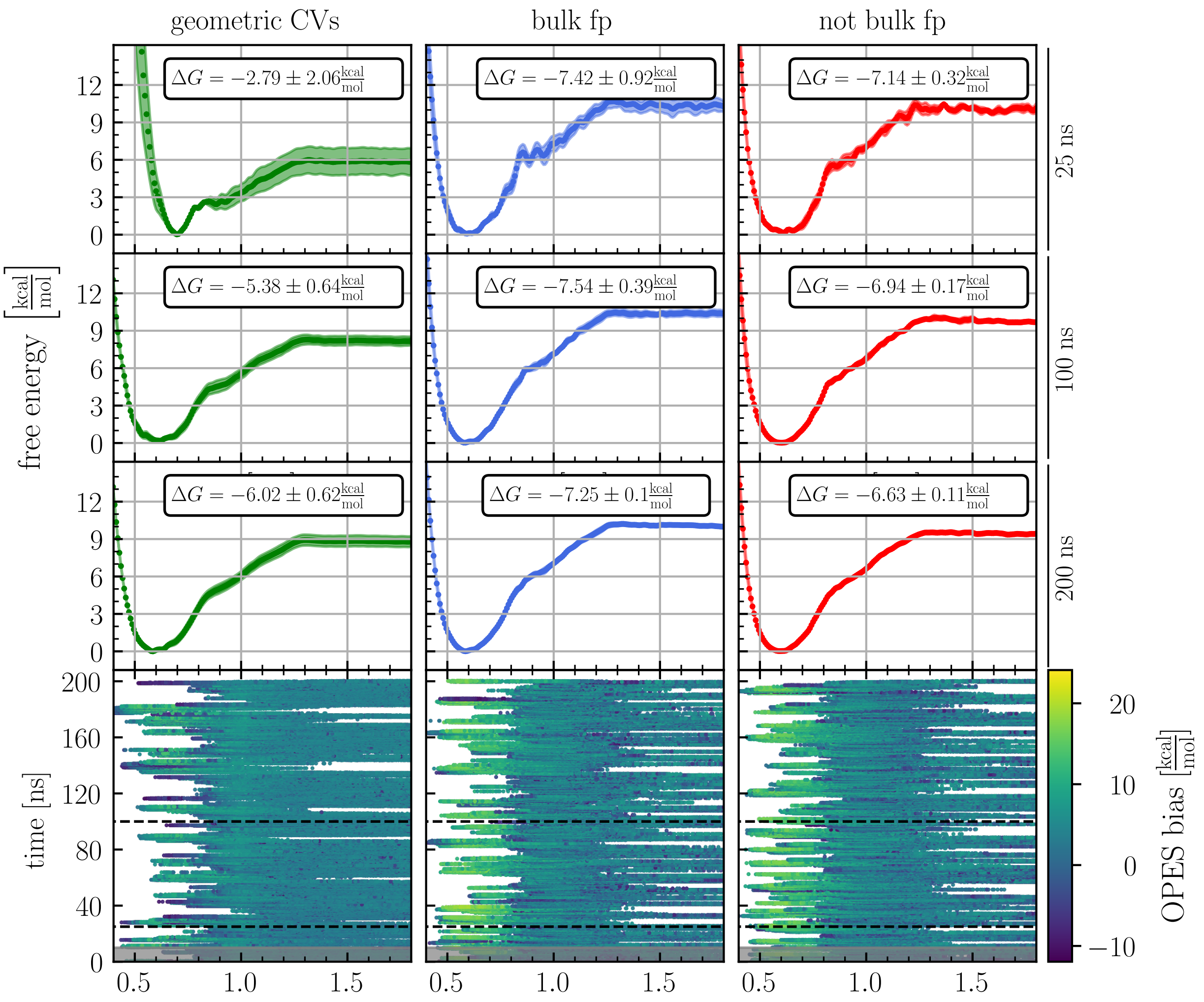


Figure S10: Free energy profiles as a function of $z$ and dynamics of $z$ as a function time for the host TEMOA and the guest S6-Go. The columns indicate the three different CVs sets. The panels in the first three rows show the averaged free energy profiles (dotted lines), their standard deviations (colored area), and the corresponding binding free energies $\Delta G$ after 25, 100, and 200 ns of sampling, respectively. The bottom row shows the time evolution of the $z$ funnel CV colored by the deposited bias. The first 10 ns shaded in gray are considered equilibration and discarded.

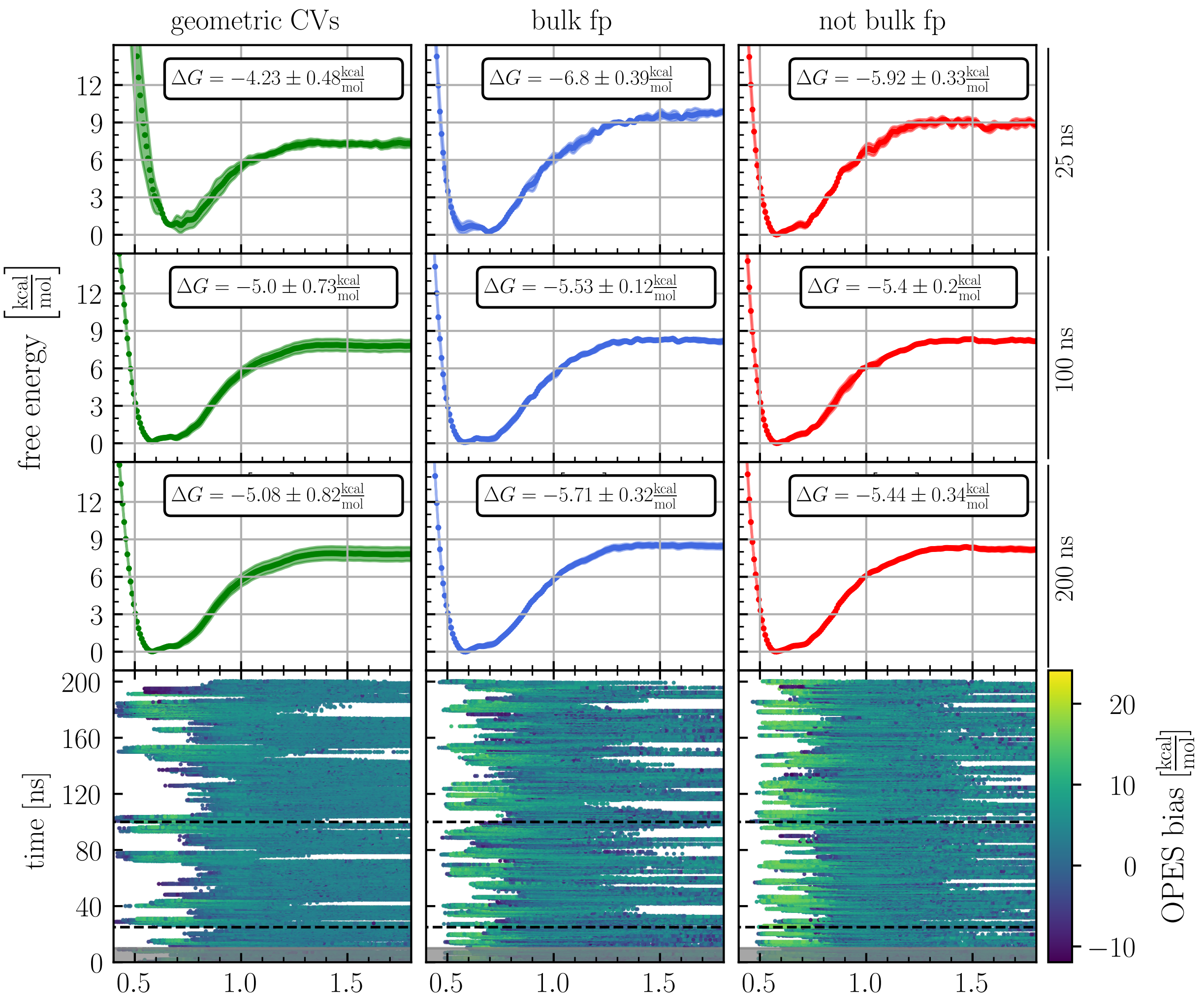


Figure S11: Free energy profiles as a function of $z$ and dynamics of $z$ as a function time for the host TEMOA and the guest S6-G1. The columns indicate the three different CVs sets. The panels in the first three rows show the averaged free energy profiles (dotted lines), their standard deviations (colored area), and the corresponding binding free energies $\Delta G$ after 25, 100, and 200 ns of sampling, respectively. The bottom row shows the time evolution of the $z$ funnel CV colored by the deposited bias. The first 10 ns shaded in gray are considered equilibration and discarded.

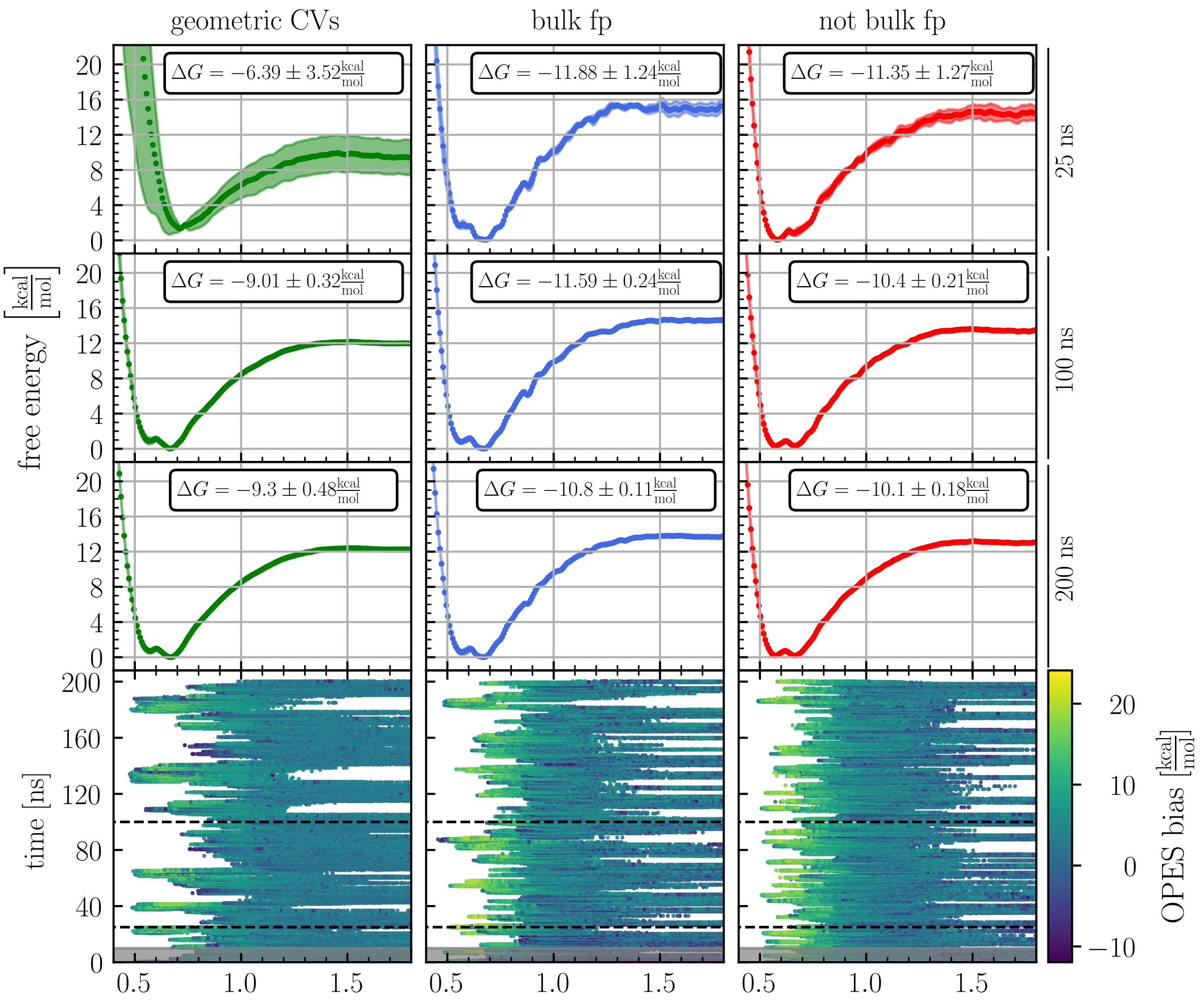


Figure S12: Free energy profiles as a function of $z$ and dynamics of $z$ as a function time for the host TEMOA and the guest S6-G2. The columns indicate the three different CVs sets. The panels in the first three rows show the averaged free energy profiles (dotted lines), their standard deviations (colored area), and the corresponding binding free energies $\Delta G$ after 25, 100, and 200 ns of sampling, respectively. The bottom row shows the time evolution of the $z$ funnel CV colored by the deposited bias. The first 10 ns shaded in gray are considered equilibration and discarded.

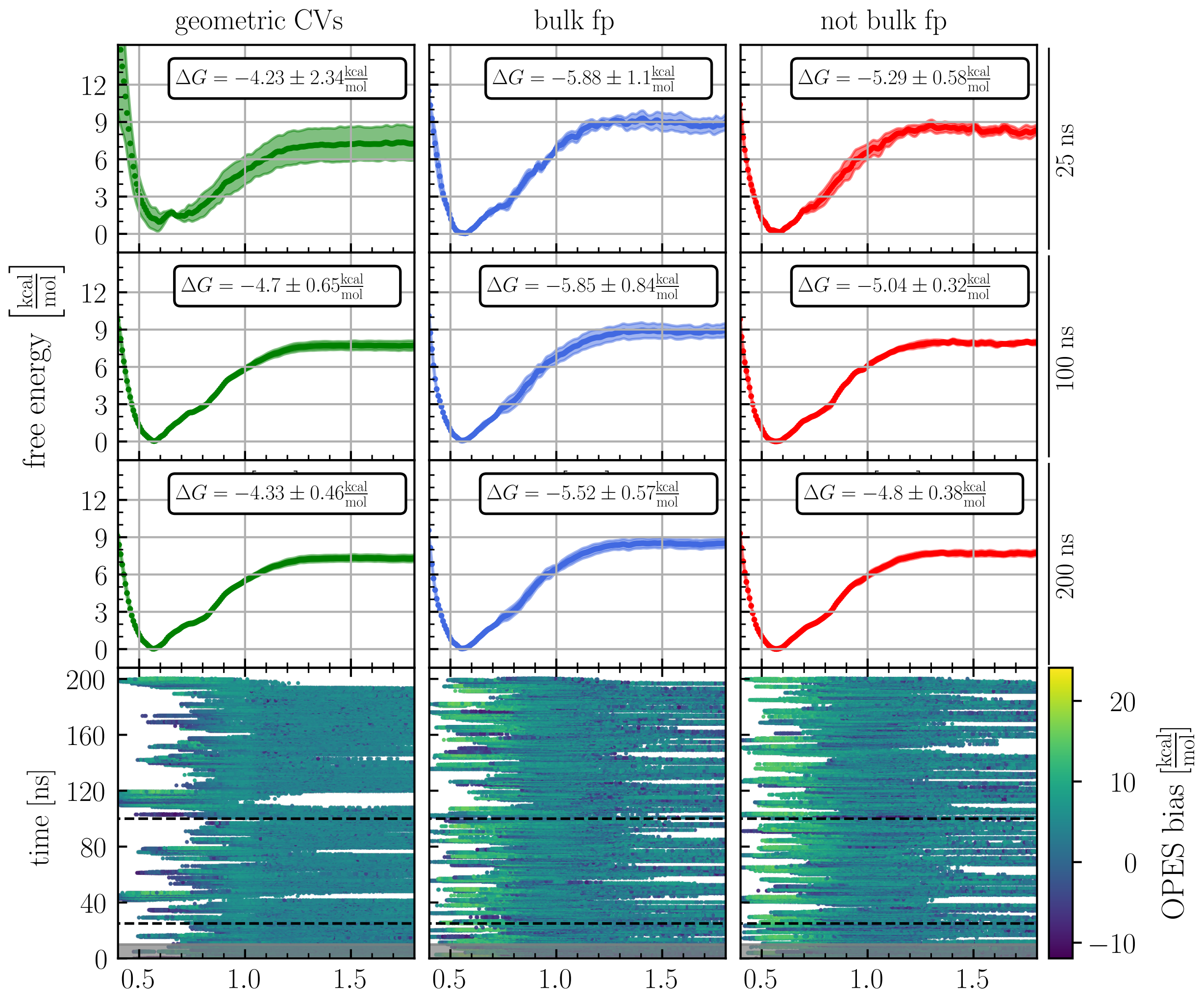


Figure S13: Free energy profiles as a function of $z$ and dynamics of $z$ as a function time for the host TEMOA and the guest S6-G3. The columns indicate the three different CVs sets. The panels in the first three rows show the averaged free energy profiles (dotted lines), their standard deviations (colored area), and the corresponding binding free energies $\Delta G$ after 25, 100, and 200 ns of sampling, respectively. The bottom row shows the time evolution of the $z$ funnel CV colored by the deposited bias. The first 10 ns shaded in gray are considered equilibration and discarded.

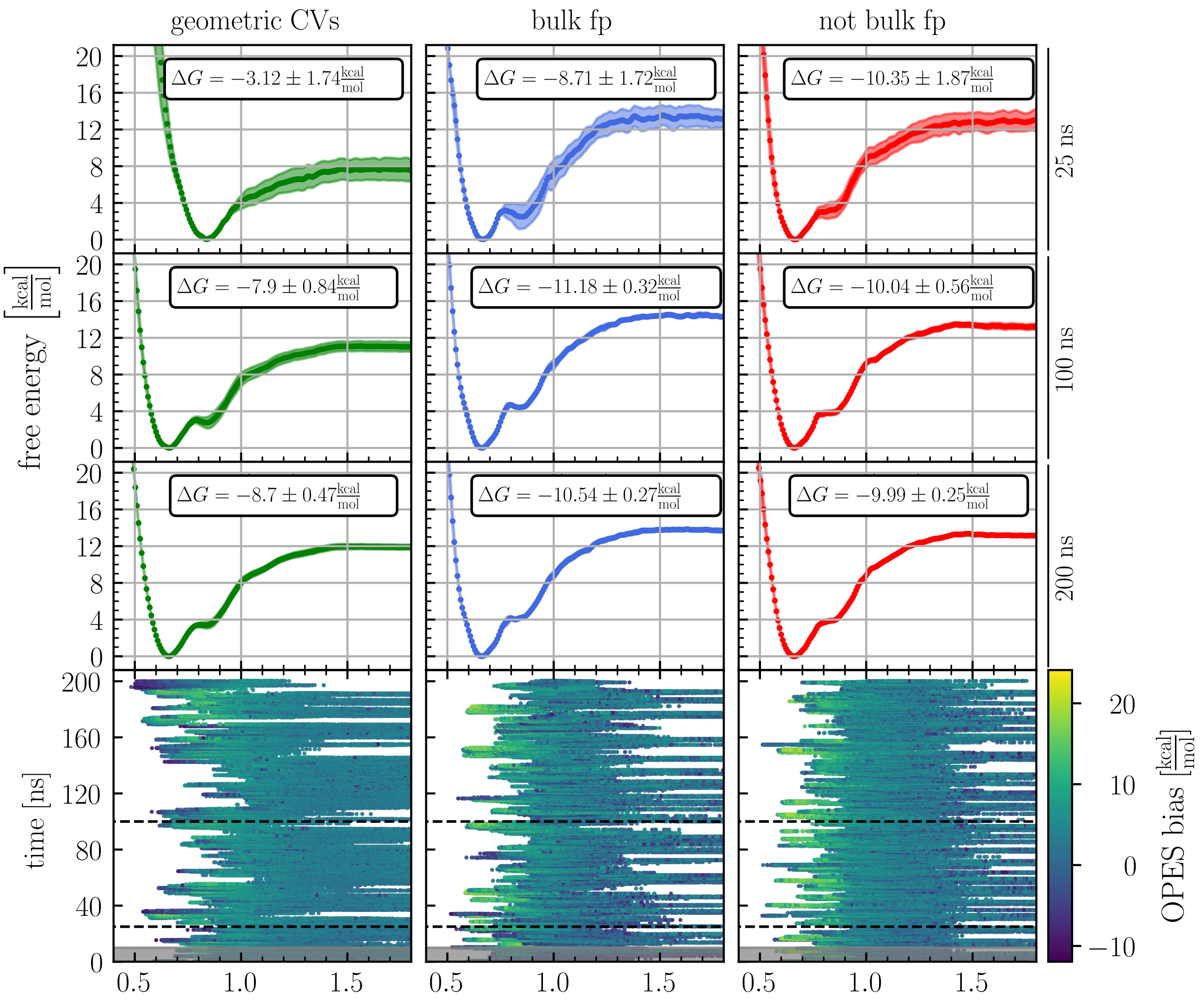


Figure S14: Free energy profiles as a function of $z$ and dynamics of $z$ as a function time for the host TEMOA and the guest S6-G4. The columns indicate the three different CVs sets. The panels in the first three rows show the averaged free energy profiles (dotted lines), their standard deviations (colored area), and the corresponding binding free energies $\Delta G$ after 25, 100, and 200 ns of sampling, respectively. The bottom row shows the time evolution of the $z$ funnel CV colored by the deposited bias. The first 10 ns shaded in gray are considered equilibration and discarded.

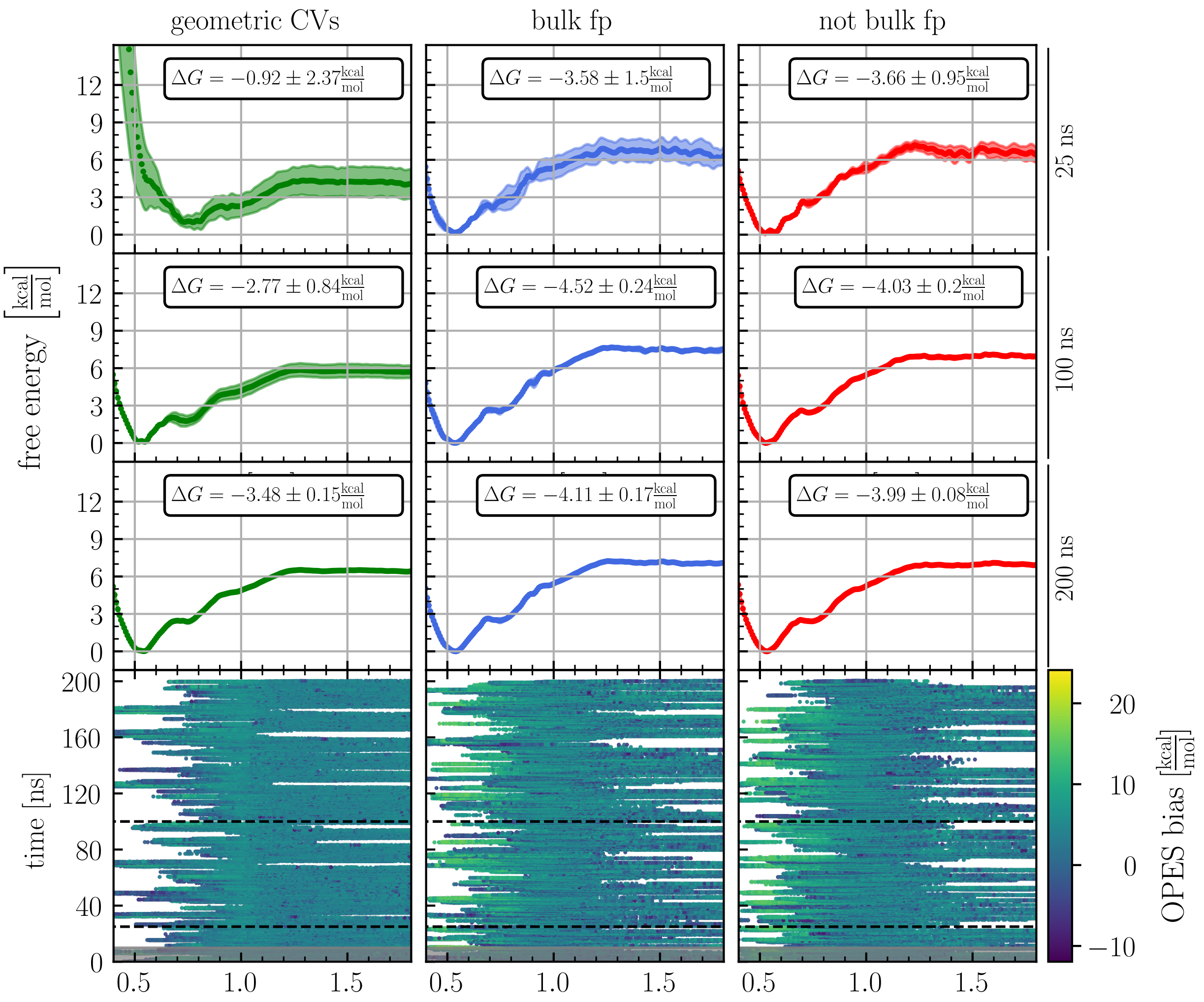


Figure S15: Free energy profiles as a function of $z$ and dynamics of $z$ as a function time for the host TEMOA and the guest S6-G5. The columns indicate the three different CVs sets. The panels in the first three rows show the averaged free energy profiles (dotted lines), their standard deviations (colored area), and the corresponding binding free energies $\Delta G$ after 25, 100, and 200 ns of sampling, respectively. The bottom row shows the time evolution of the $z$ funnel CV colored by the deposited bias. The first 10 ns shaded in gray are considered equilibration and discarded.

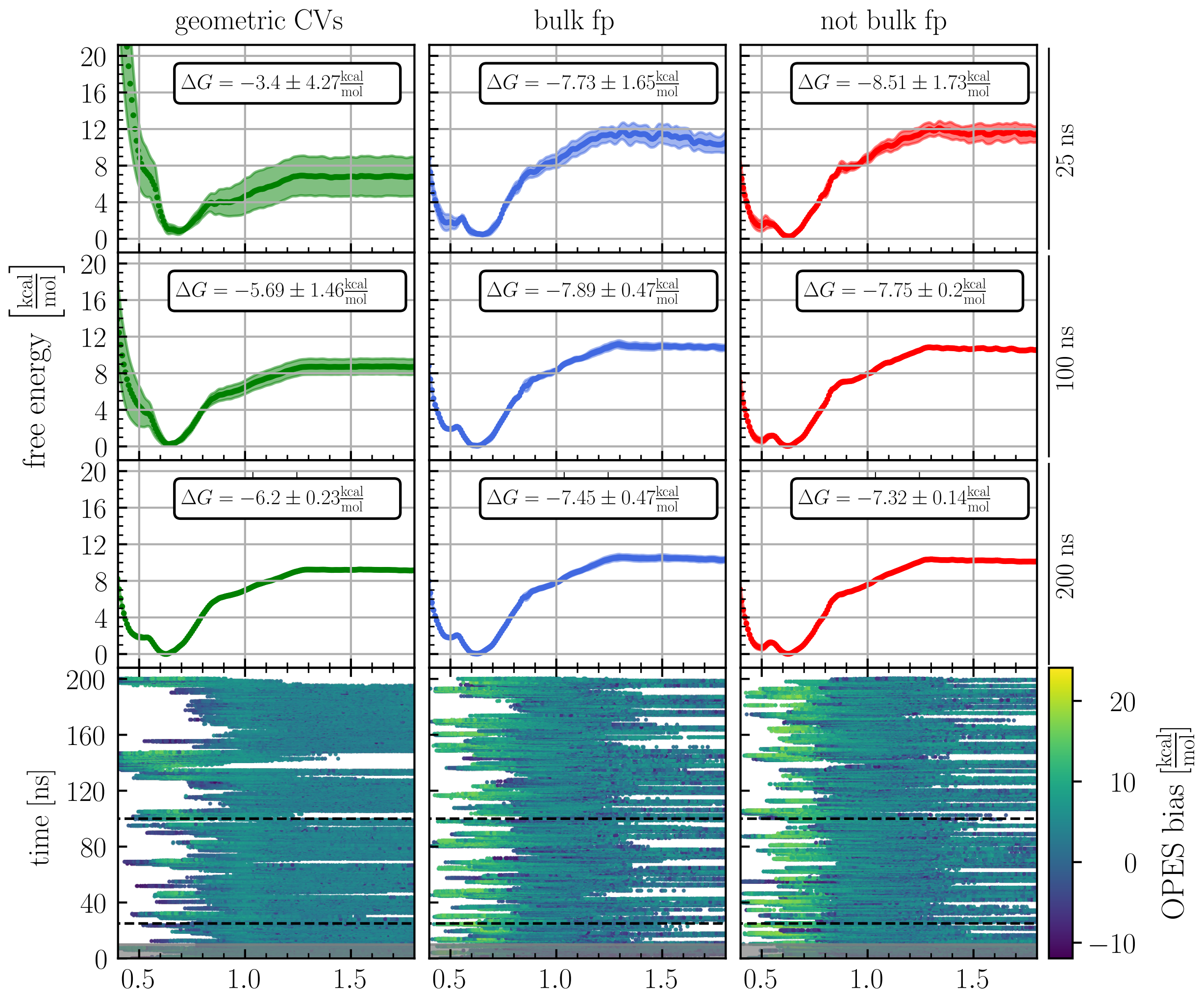


Figure S16: Free energy profiles as a function of $z$ and dynamics of $z$ as a function time for the host TEMOA and the guest S6-G6. The columns indicate the three different CVs sets. The panels in the first three rows show the averaged free energy profiles (dotted lines), their standard deviations (colored area), and the corresponding binding free energies $\Delta G$ after 25, 100, and 200 ns of sampling, respectively. The bottom row shows the time evolution of the $z$ funnel CV colored by the deposited bias. The first 10 ns shaded in gray are considered equilibration and discarded.

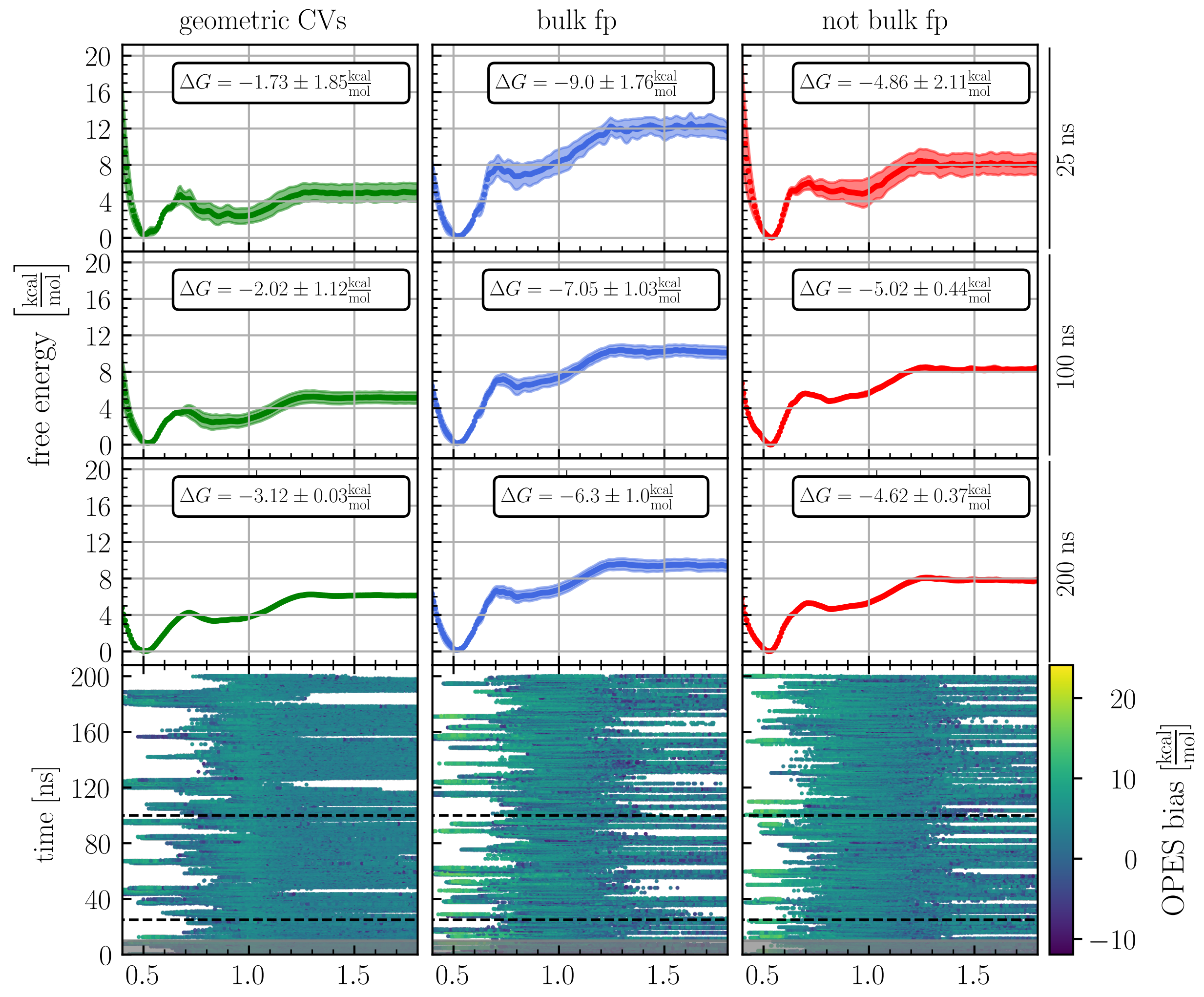


Figure S17: Free energy profiles as a function of $z$ and dynamics of $z$ as a function time for the host TEMOA and the guest S6-G7. The columns indicate the three different CVs sets. The panels in the first three rows show the averaged free energy profiles (dotted lines), their standard deviations (colored area), and the corresponding binding free energies $\Delta G$ after 25, 100, and 200 ns of sampling, respectively. The bottom row shows the time evolution of the $z$ funnel CV colored by the deposited bias. The first 10 ns shaded in gray are considered equilibration and discarded.

### Statistical estimators for all the CV sets

In this section, we report in Figures S18 and S19 a direct comparison of the performances of our OPES simulations carried out with the three sets of CVs, i.e., geometrical (green), bulk (blue), and anti-bulk (red).

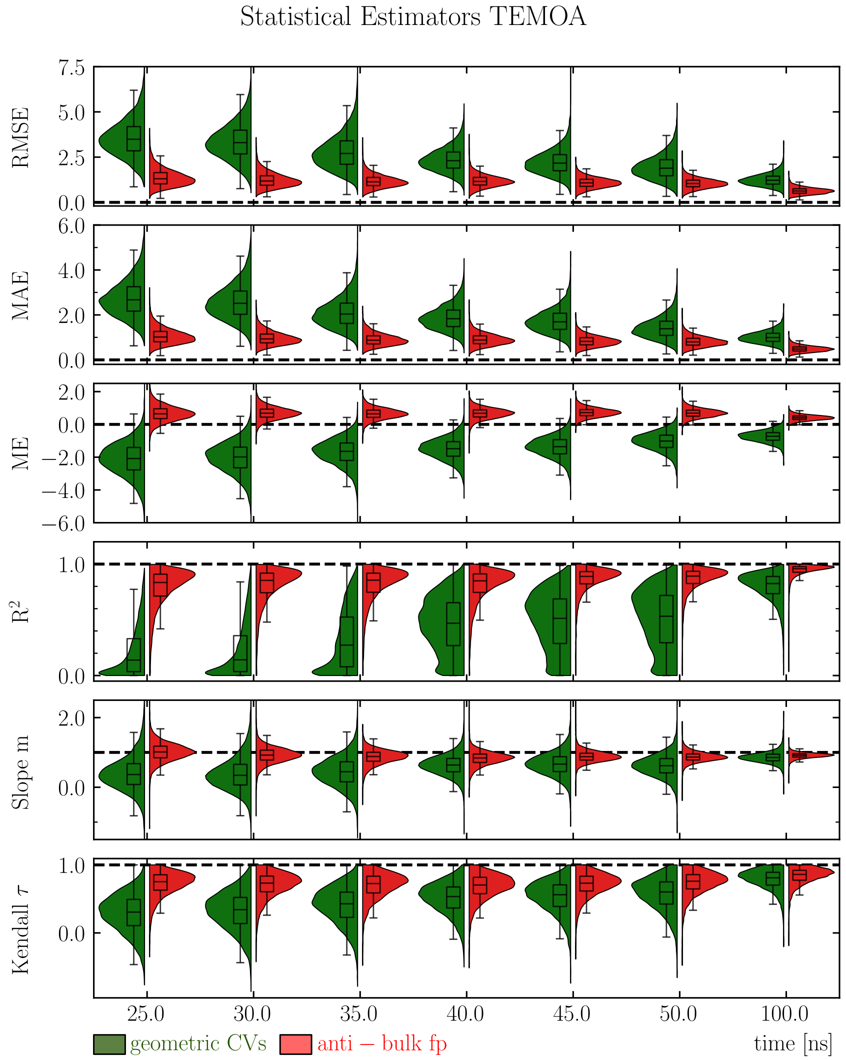


Figure S18: Time evolution of statistical estimators for the geometrical (green) and anti-bulk fingerprint (red) CVs sets. In addition to the $R^2$ and RMSE reported also in the main text, we computed the mean absolute error (MAE), the mean error (ME), the fitted slope of the estimated vs. reference $\Delta G$ (Slope $m$), and the Kendall rank correlation coefficient (Kendall $\tau$). The confidence intervals are calculated by bootstrapping.

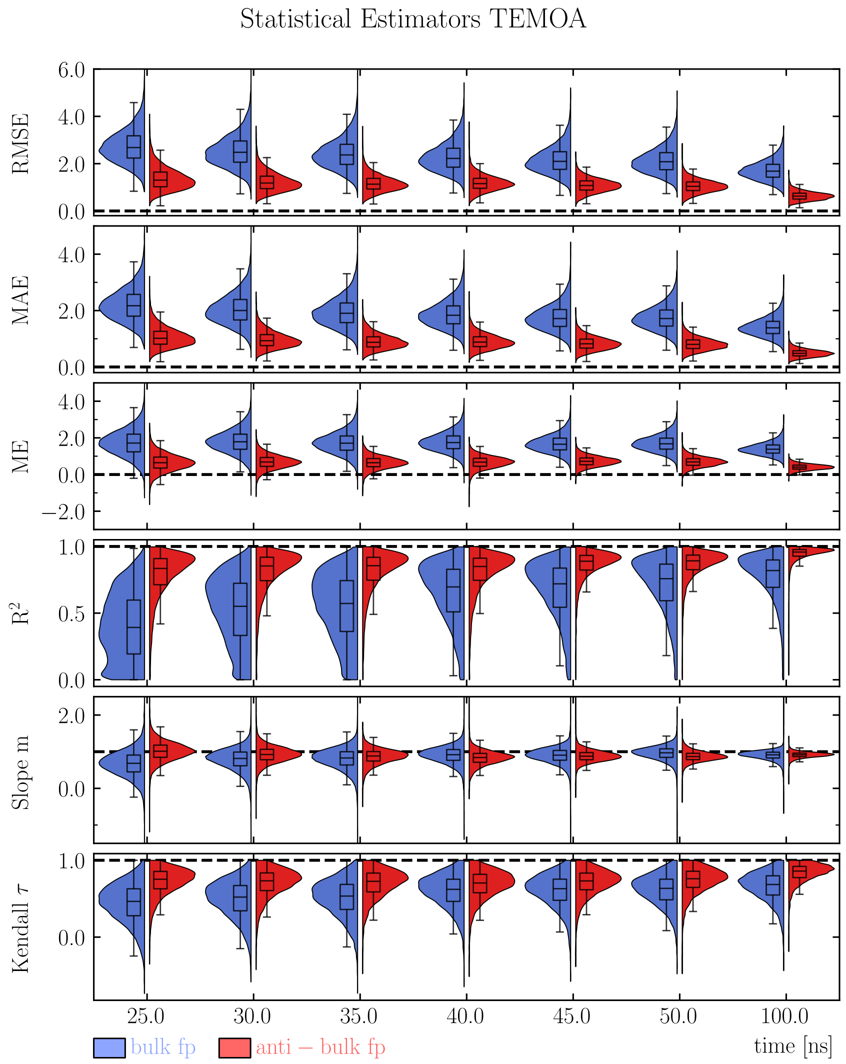


Figure S19: Time evolution of statistical estimators for the bulk (blue) and anti-bulk (red) fingerprint CVs sets. In addition to the $R^2$ and RMSE reported also in the main text, we computed the mean absolute error (MAE), the mean error (ME), the fitted slope of the estimated vs. reference $\Delta G$ (Slope $m$), and the Kendall rank correlation coefficient (Kendall $\tau$). The confidence intervals are calculated by bootstrapping.

### Free energy convergence at long time scales

The (long) time evolution at 200 ns and 500 ns of the binding free-energy estimates was monitored for the host-guest complexes studied in this work. In Figure S20 the estimates are compared against reference values to assess convergence and stability. These plots illustrate how the calculated free energies approach the reference values over simulation time regardless of the set of CVs employed, thereby validating the reliability of our enhanced sampling protocol.

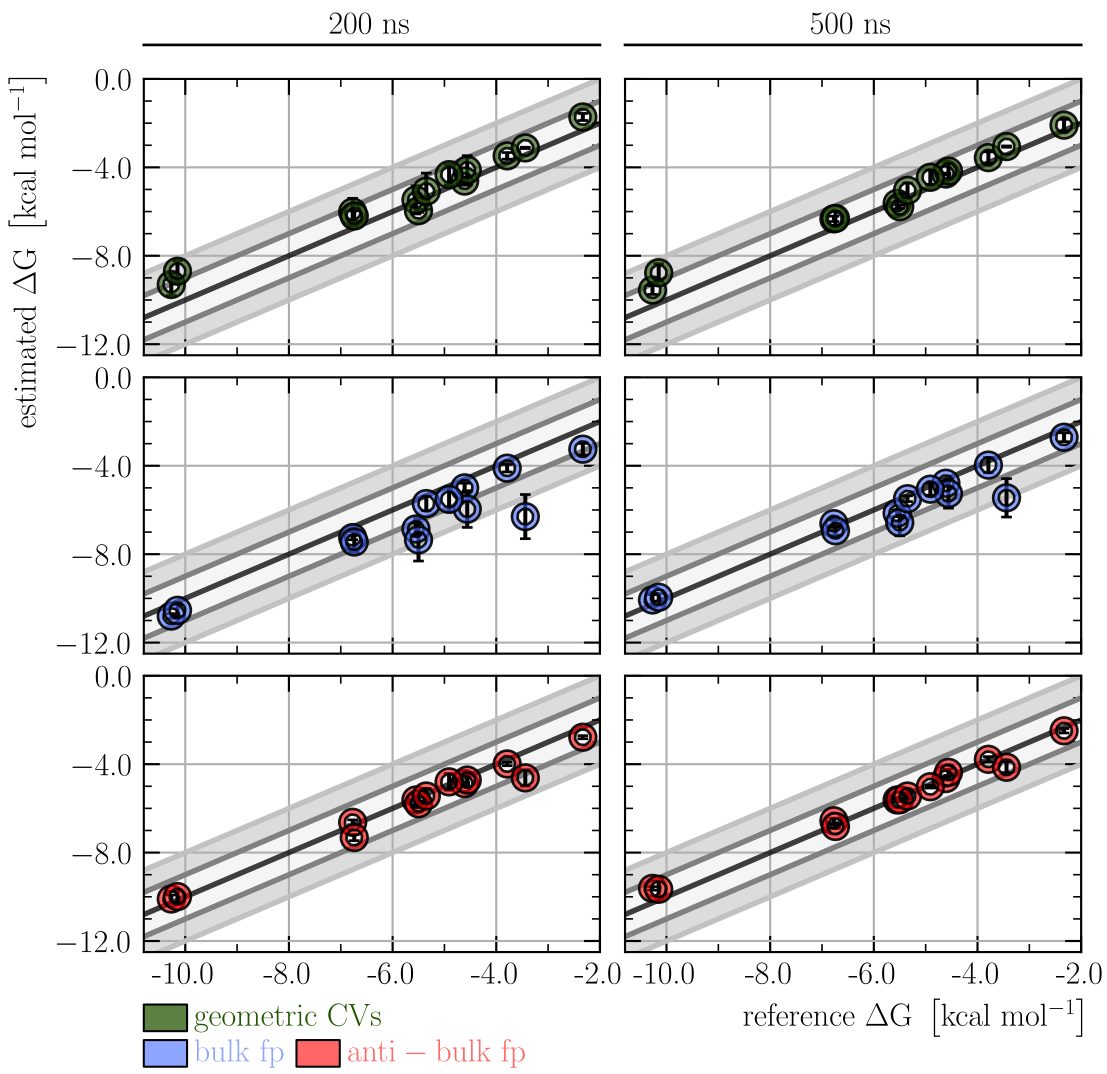


Figure S20: Comparison of the free energy estimates after 200 (left) and 500 (right) ns of sampling to the reference values for all the CVs sets. Average and one standard deviation of three independent simulations.

### Free energy differences over time for all ligands and CV sets

Here we report the time evolution of ΔG of binding for the host-guest complexes involved in SAMPL5 (see Figure S21) and SAMPL6 (see Figure S22).

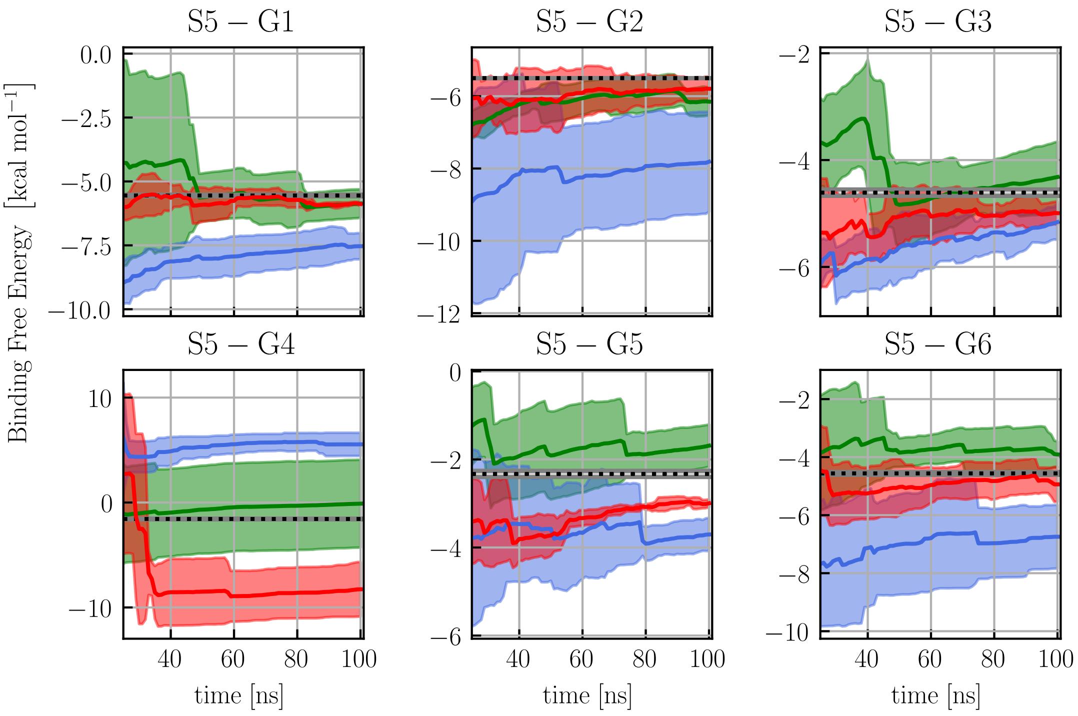


Figure S21: Free energy estimates at different simulation lengths for the three CVs sets for all the guest of SAMPL5. Average and standard deviation of three independent simulations. The corresponding references are dotted black lines.

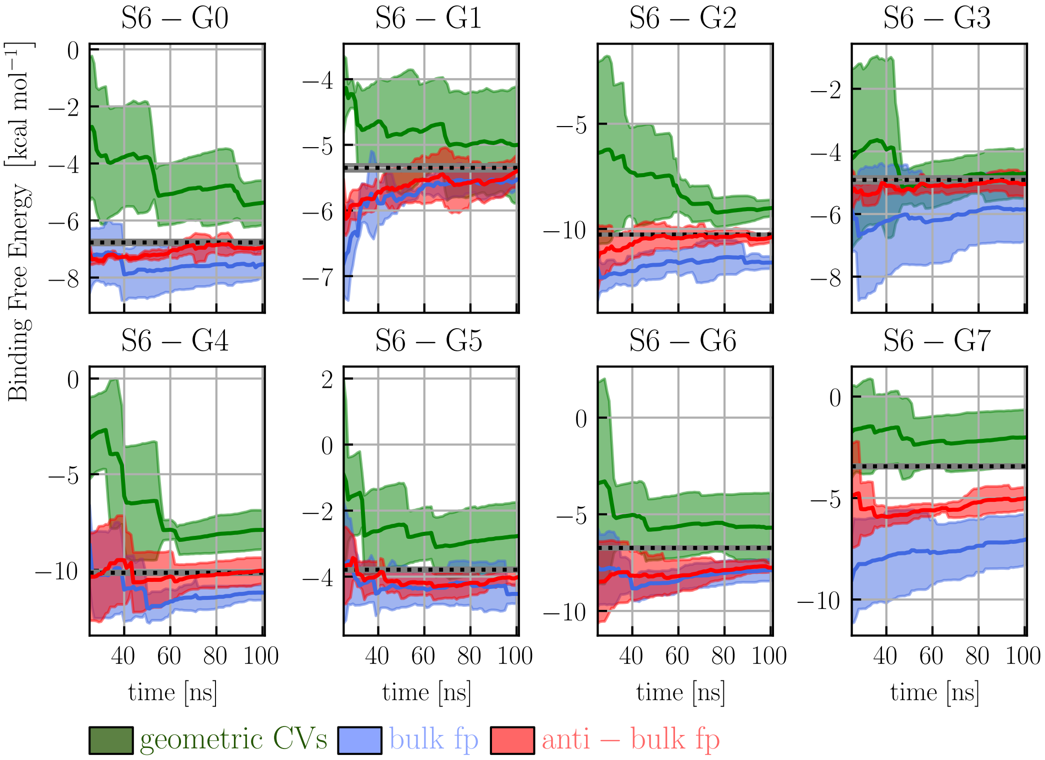


Figure S22: Free energy estimates at different simulation lengths for the three CVs sets for all the guest of SAMPL6. Average and standard deviation of three independent simulations. The corresponding references are dotted black lines.

### Comparison with experimental data

In this section we report the computed binding free energies for all host–guest systems and compare them with the corresponding experimental reference values. Results for the three sets of CVs are shown in Figure S23, Figure S24, and Figure S25, enabling a direct assessment of the accuracy and consistency of the calculations across different biasing schemes.

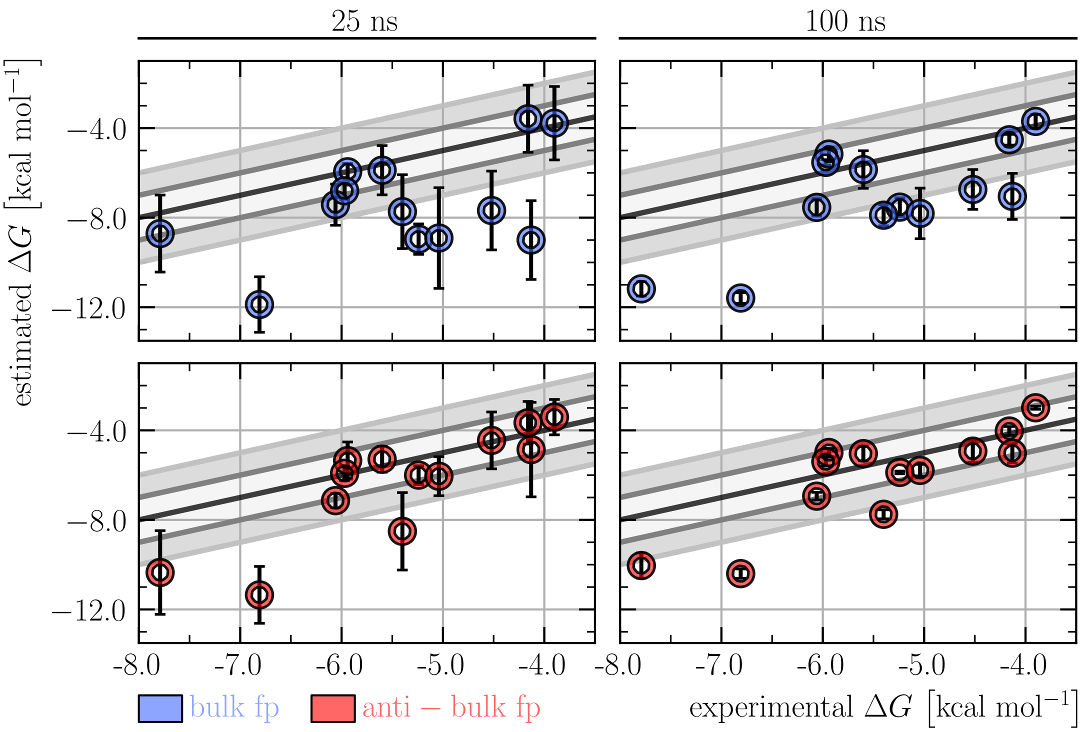


Figure S23: Comparison of the free energy estimates after 25 (left) and 100 (right) ns of sampling to the experimental values for bulk (blue) and anti-bulk (red) water CVs sets. Average and one standard deviation of three independent simulations.

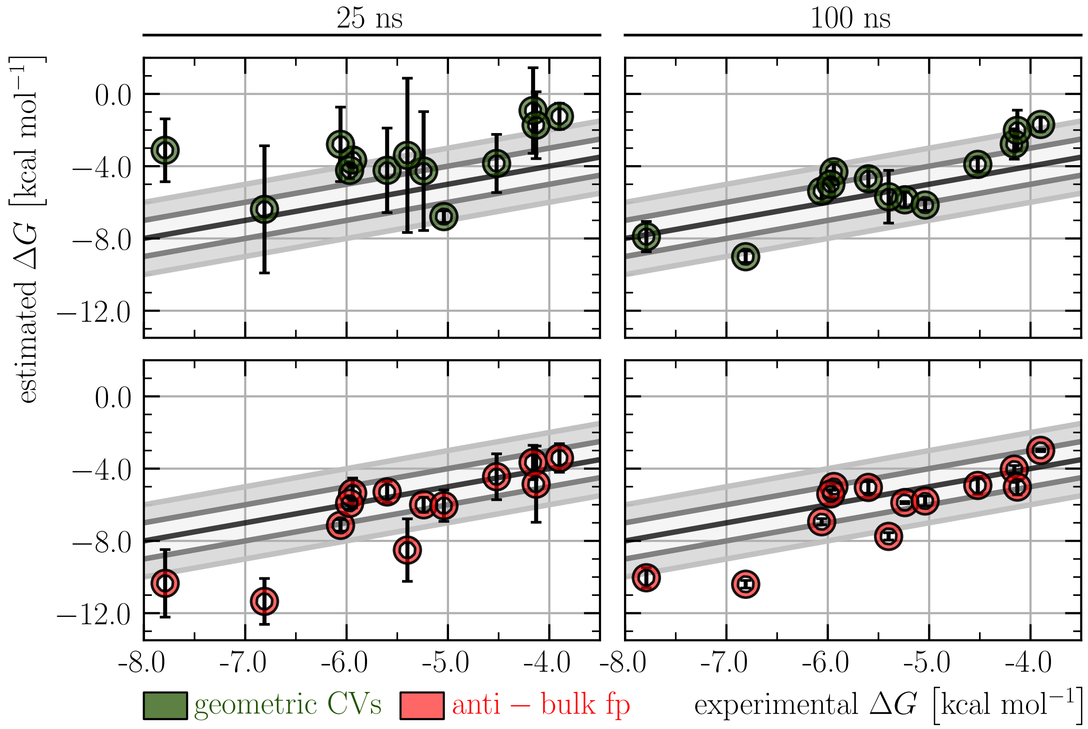


Figure S24: Comparison of the free energy estimates after 25 (left) and 100 (right) ns of sampling to the experimental values for anti-bulk (blue) and geometric (green) CVs sets. Average and one standard deviation of three independent simulations.

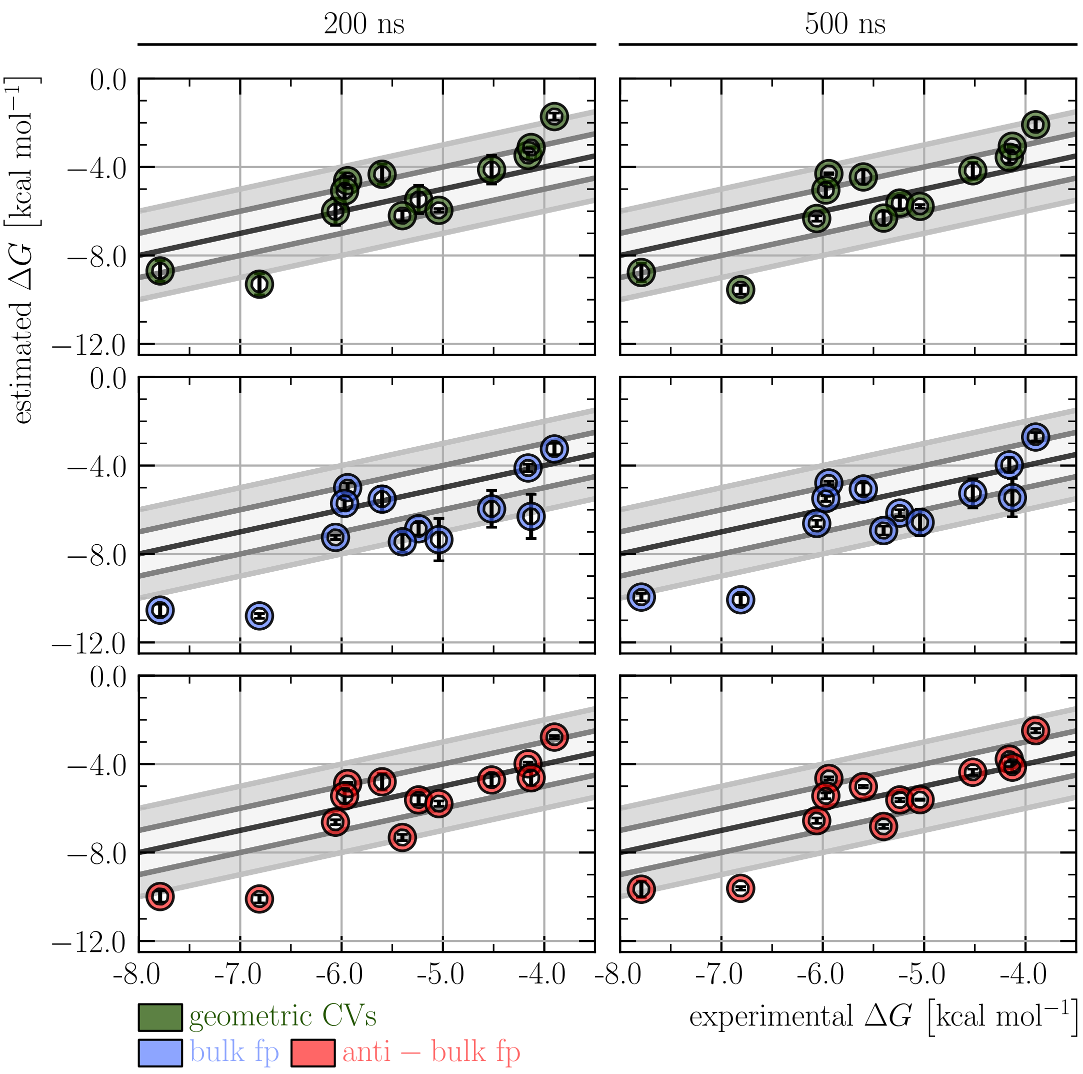


Figure S25: Comparison of the free energy estimates after 200 (left) and 500 (right) ns of sampling to the experimental values for all three CV sets. Average and one standard deviation of three independent simulations.

### Solvent accessible surface area analysis

Solvent Accessible Surface Area (SASA) values of all heavy atoms of each ligand were calculated in VMD (Visual Molecular Dynamics) [1], using a radius of 1.4 Å. These values were computed from the same trajectories that were used to calculate the fingerprint. The SASA values of the ligand atoms were then compared to their corresponding fingerprint by computing both the Spearman rank correlation and the Pearson coefficients to assess monotonic and linear relationships, respectively. The correlation coefficients were computed using the scipy.stats Python module [2].

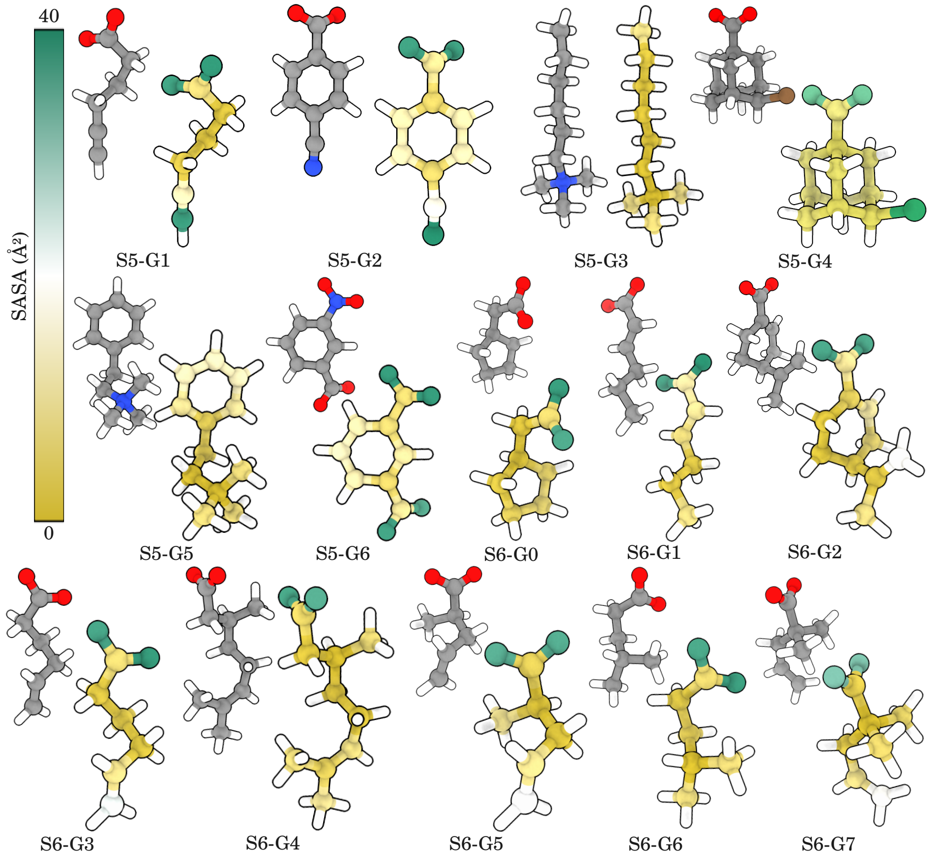


Figure S26: Global comparison of solvent accessible surface area (SASA) values of SAMPL5 and SAMPL6 guest molecules. Each guest molecule is shown in two representations. On the left, its atoms are colored according to their element type (oxygen, nitrogen, carbon, and hydrogen atoms are colored in red, blue, gray and white, respectively). On the right, each atom of a guest molecules is colored according to its SASA value, except for hydrogen ones, which are only shown for clarity.

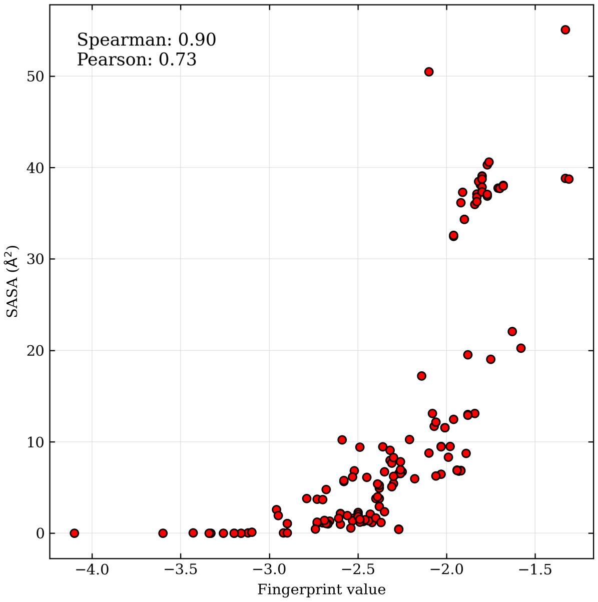


Figure S27: Correlation between fingerprint values and solvent-accessible surface area (SASA) across all heavy atoms of ligand systems.

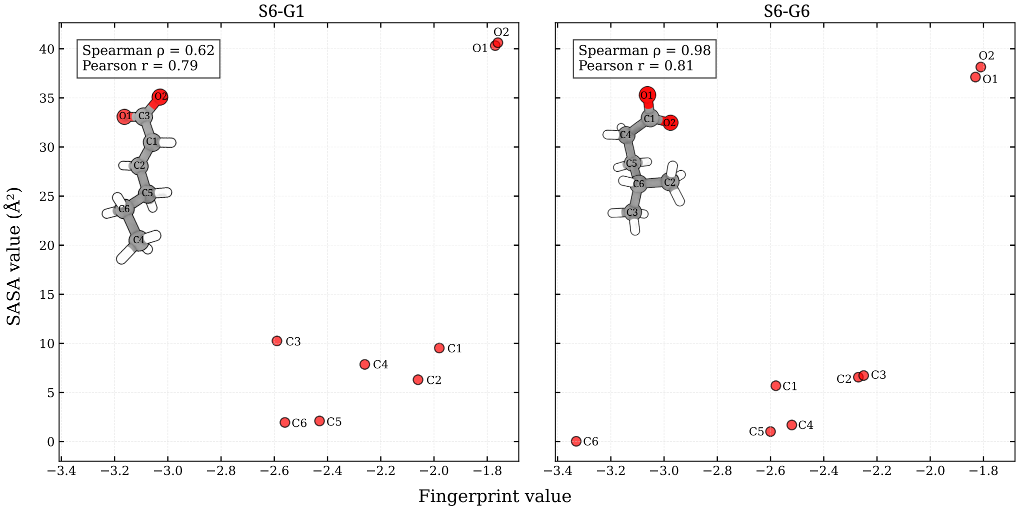


Figure S28: Correlation between fingerprint values and solvent accessible surface area (SASA) for heavy atom of ligand S6-G1 and S6-G6. The ligand representation is colored according to the element type (oxygen in red, carbon in gray, and hydrogen in white) and the atom labels correspond to the ones in the plot.

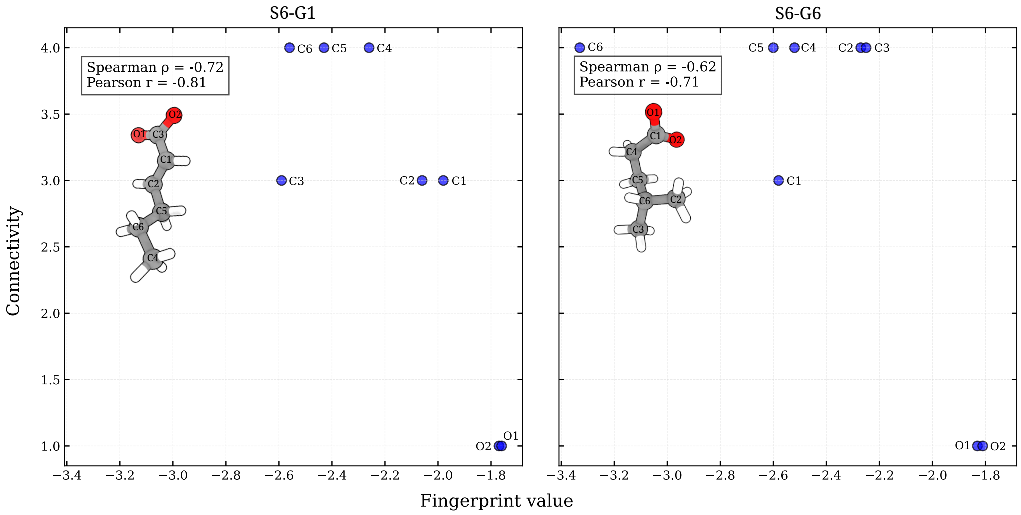


Figure S29: Correlation between fingerprint values and atom connectivity for all heavy atoms of ligand S6-G1 and S6-G6. The ligand representation is colored according to the element type (oxygen in red, carbon in gray, and hydrogen in white) and the atom labels correspond to the ones in the plot.

### Predicted ΔG values

In this section, we explicitly report the ΔG values and their relative errors (one standard deviation).

| Host-Guest | $\Delta G_{\text{reference}}$ $\left[\frac{\text{kcal}}{\text{mol}}\right]$ | $\Delta G^{\text{estimated}}_{\text{(geometric CVs)}}$ $\left[\frac{\text{kcal}}{\text{mol}}\right]$ | $\Delta G^{\text{estimated}}_{\text{(bulk)}}$ $\left[\frac{\text{kcal}}{\text{mol}}\right]$ | $\Delta G^{\text{estimated}}_{\text{(anti-bulk)}}$ $\left[\frac{\text{kcal}}{\text{mol}}\right]$ |
|---|---|---|---|---|
| TEMOA-S5-G1 | $-5.55 \pm 0.03$ | $-4.27 \pm 3.29$ | $-8.96 \pm 0.68$ | $-6.00 \pm 0.42$ |
| TEMOA-S5-G2 | $-5.50 \pm 0.00$ | $-6.79 \pm 0.32$ | $-8.91 \pm 2.25$ | $-6.05 \pm 0.87$ |
| TEMOA-S5-G3 | $-4.61 \pm 0.06$ | $-3.66 \pm 0.68$ | $-5.95 \pm 0.37$ | $-5.35 \pm 0.83$ |
| TEMOA-S5-G4* | $-1.57 \pm 0.03$ | $-1.07 \pm 3.87$ | $6.16 \pm 4.33$ | $2.71 \pm 6.21$ |
| TEMOA-S5-G5 | $-2.33 \pm 0.07$ | $-1.24 \pm 0.72$ | $-3.78 \pm 1.64$ | $-3.41 \pm 0.79$ |
| TEMOA-S5-G6 | $-4.56 \pm 0.05$ | $-3.85 \pm 1.61$ | $-7.68 \pm 1.76$ | $-4.45 \pm 1.27$ |
| TEMOA-S6-G0 | $-6.77 \pm 0.06$ | $-2.79 \pm 2.06$ | $-7.42 \pm 0.92$ | $-7.14 \pm 0.32$ |
| TEMOA-S6-G1 | $-5.35 \pm 0.04$ | $-4.23 \pm 0.48$ | $-6.80 \pm 0.39$ | $-5.92 \pm 0.33$ |
| TEMOA-S6-G2 | $-10.27 \pm 0.00$ | $-6.39 \pm 3.52$ | $-11.88 \pm 1.24$ | $-11.35 \pm 1.27$ |
| TEMOA-S6-G3 | $-4.91 \pm 0.08$ | $-4.23 \pm 2.34$ | $-5.88 \pm 1.10$ | $-5.29 \pm 0.58$ |
| TEMOA-S6-G4 | $-10.15 \pm 0.06$ | $-3.12 \pm 1.74$ | $-8.71 \pm 1.72$ | $-10.35 \pm 1.87$ |
| TEMOA-S6-G5 | $-3.79 \pm 0.03$ | $-0.92 \pm 2.37$ | $-3.58 \pm 1.50$ | $-3.66 \pm 0.95$ |
| TEMOA-S6-G6 | $-6.74 \pm 0.03$ | $-3.40 \pm 4.27$ | $-7.73 \pm 1.65$ | $-8.51 \pm 1.73$ |
| TEMOA-S6-G7 | $-3.44 \pm 0.05$ | $-1.73 \pm 1.85$ | $-9.00 \pm 1.76$ | $-4.86 \pm 2.11$ |

Table S1: Free energy estimates for host-guest systems at 25 ns of simulation for all the different CV sets. One standard deviation shown as ±. Guest S5-G4 (*) has been removed from the results presented in the main text as the hydration CV sets cannot distinguish between the intermediate and the bound state.

| Host-Guest | $\Delta G_{\text{reference}}$ $\left[\frac{\text{kcal}}{\text{mol}}\right]$ | $\Delta G^{\text{estimated}}_{\text{(geometric CVs)}}$ $\left[\frac{\text{kcal}}{\text{mol}}\right]$ | $\Delta G^{\text{estimated}}_{\text{(bulk)}}$ $\left[\frac{\text{kcal}}{\text{mol}}\right]$ | $\Delta G^{\text{estimated}}_{\text{(anti-bulk)}}$ $\left[\frac{\text{kcal}}{\text{mol}}\right]$ |
|---|---|---|---|---|
| TEMOA-S5-G1 | $-5.55 \pm 0.03$ | $-5.86 \pm 0.45$ | $-7.53 \pm 0.42$ | $-5.88 \pm 0.06$ |
| TEMOA-S5-G2 | $-5.50 \pm 0.00$ | $-6.15 \pm 0.30$ | $-7.81 \pm 1.13$ | $-5.79 \pm 0.27$ |
| TEMOA-S5-G3 | $-4.61 \pm 0.06$ | $-4.31 \pm 0.54$ | $-5.16 \pm 0.27$ | $-4.99 \pm 0.18$ |
| TEMOA-S5-G4* | $-1.57 \pm 0.03$ | $-0.11 \pm 3.42$ | $5.54 \pm 0.91$ | $-8.26 \pm 2.12$ |
| TEMOA-S5-G5 | $-2.33 \pm 0.07$ | $-1.69 \pm 0.39$ | $-3.70 \pm 0.31$ | $-2.99 \pm 0.07$ |
| TEMOA-S5-G6 | $-4.56 \pm 0.05$ | $-3.90 \pm 0.37$ | $-6.74 \pm 0.89$ | $-4.93 \pm 0.49$ |
| TEMOA-S6-G0 | $-6.77 \pm 0.06$ | $-5.38 \pm 0.64$ | $-7.54 \pm 0.39$ | $-6.94 \pm 0.17$ |
| TEMOA-S6-G1 | $-5.35 \pm 0.04$ | $-5.00 \pm 0.73$ | $-5.53 \pm 0.12$ | $-5.40 \pm 0.20$ |
| TEMOA-S6-G2 | $-10.27 \pm 0.00$ | $-9.01 \pm 0.32$ | $-11.59 \pm 0.24$ | $-10.40 \pm 0.21$ |
| TEMOA-S6-G3 | $-4.91 \pm 0.08$ | $-4.70 \pm 0.65$ | $-5.85 \pm 0.84$ | $-5.04 \pm 0.32$ |
| TEMOA-S6-G4 | $-10.15 \pm 0.06$ | $-7.90 \pm 0.84$ | $-11.18 \pm 0.32$ | $-10.04 \pm 0.56$ |
| TEMOA-S6-G5 | $-3.79 \pm 0.03$ | $-2.77 \pm 0.84$ | $-4.52 \pm 0.24$ | $-4.03 \pm 0.20$ |
| TEMOA-S6-G6 | $-6.74 \pm 0.03$ | $-5.69 \pm 1.46$ | $-7.89 \pm 0.47$ | $-7.75 \pm 0.20$ |
| TEMOA-S6-G7 | $-3.44 \pm 0.05$ | $-2.02 \pm 1.12$ | $-7.05 \pm 1.03$ | $-5.02 \pm 0.44$ |

Table S2: Free energy estimates for host-guest systems at 100 ns of simulation for all the different CV sets. One standard deviation shown as ±. Guest S5-G4 (*) has been removed from the results presented in the main text as the hydration CV sets cannot distinguish between the intermediate and the bound state.

### Predicted fingerprint values for different water models

In this section, we list the fingerprint values we measured for the atoms of all SAMPL5 and SAMPL6 guests for the TIP3P water model (employed in the main text) and three other different water models (SPC, TIP4P, and TIP4Pew). It can be seen how for all ligands the ordering of the atoms depends very weakly on the water model as the classification is consistent among models and, when not, it results in a switch of consecutive positions that depends on the second significative digit of the fingerprint. Nevertheless, to ensure consistency, we recommend recalculating the fingerprint values if the water model or the employed force field change.

| Ligand | TIP3P | | SPC | | TIP4P | | TIP4Pew | |
|---|---|---|---|---|---|---|---|---|
| | Atom | Fingerprint | Atom | Fingerprint | Atom | Fingerprint | Atom | Fingerprint |
| | 6 | -2,52 | 6 | -2,82 | 6 | -2,67 | 6 | -2,79 |
| | 5 | -2,46 | 5 | -2,66 | 5 | -2,63 | 5 | -2,70 |
| | 4 | -2,44 | 4 | -2,66 | 4 | -2,60 | 4 | -2,66 |
| S5-G1 | 3 | -2,35 | 3 | -2,57 | 3 | -2,52 | 3 | -2,57 |
| | 2 | -1,96 | 2 | -2,16 | 2 | -2,10 | 2 | -2,14 |
| | 7 | -1,80 | 8 | -2,05 | 7 | -1,96 | 7 | -2,10 |
| | 1 | -1,80 | 7 | -2,05 | 8 | -1,96 | 8 | -2,08 |
| | 8 | -1,80 | 1 | -1,99 | 1 | -1,94 | 1 | -1,99 |
| | 3 | -2,39 | 9 | -2,53 | 3 | -2,52 | 9 | -2,58 |
| | 9 | -2,32 | 3 | -2,53 | 9 | -2,51 | 3 | -2,57 |
| | 6 | -2,31 | 6 | -2,44 | 6 | -2,46 | 6 | -2,53 |
| | 7 | -2,14 | 7 | -2,27 | 7 | -2,27 | 7 | -2,33 |
| | 4 | -2,06 | 4 | -2,19 | 4 | -2,19 | 4 | -2,24 |
| S5-G2 | 2 | -2,06 | 2 | -2,19 | 2 | -2,19 | 2 | -2,24 |
| | 1 | -2,01 | 5 | -2,16 | 1 | -2,16 | 5 | -2,21 |
| | 5 | -2,01 | 1 | -2,16 | 5 | -2,16 | 1 | -2,21 |
| | 11 | -1,68 | 10 | -1,90 | 10 | -1,89 | 11 | -2,00 |
| | 10 | -1,68 | 11 | -1,89 | 11 | -1,89 | 10 | -1,98 |
| | 8 | -1,33 | 8 | -1,46 | 8 | -1,44 | 8 | -1,47 |
| | 7 | -3,26 | 7 | -3,46 | 7 | -3,38 | 7 | -3,40 |
| | 4 | -2,68 | 4 | -2,82 | 4 | -2,89 | 4 | -2,87 |
| | 3 | -2,67 | 3 | -2,79 | 3 | -2,87 | 3 | -2,85 |
| | 2 | -2,61 | 2 | -2,76 | 2 | -2,83 | 2 | -2,83 |
| S5-G3 | 5 | -2,54 | 5 | -2,68 | 5 | -2,74 | 5 | -2,71 |
| | 1 | -2,32 | 1 | -2,49 | 1 | -2,57 | 1 | -2,60 |
| | 6 | -2,27 | 6 | -2,41 | 6 | -2,50 | 6 | -2,47 |
| | 9 | -1,94 | 10 | -2,07 | 8 | -2,13 | 8 | -2,10 |
| | 8 | -1,94 | 9 | -2,07 | 9 | -2,13 | 9 | -2,10 |
| | 10 | -1,94 | 8 | -2,07 | 10 | -2,12 | 10 | -2,10 |
| | 6 | -3,68 | 6 | -3,77 | 6 | -3,79 | 6 | -3,90 |
| | 3 | -3,28 | 3 | -3,37 | 10 | -3,39 | 10 | -3,45 |
| | 10 | -3,27 | 10 | -3,37 | 3 | -3,39 | 3 | -3,44 |
| | 9 | -3,18 | 9 | -3,27 | 9 | -3,29 | 9 | -3,33 |
| | 2 | -2,99 | 2 | -3,09 | 2 | -3,11 | 7 | -3,17 |
| | 7 | -2,94 | 7 | -3,05 | 4 | -3,08 | 2 | -3,17 |
| S5-G4 | 4 | -2,93 | 4 | -3,04 | 7 | -3,08 | 4 | -3,16 |
| | 11 | -2,71 | 11 | -3,04 | 11 | -2,95 | 11 | -3,09 |
| | 5 | -2,71 | 5 | -2,81 | 5 | -2,83 | 5 | -2,91 |
| | 8 | -2,68 | 8 | -2,78 | 8 | -2,81 | 8 | -2,86 |
| | 1 | -2,68 | 1 | -2,77 | 1 | -2,81 | 1 | -2,86 |
| | 14 | -2,19 | 14 | -2,30 | 14 | -2,35 | 14 | -2,44 |
| | 12 | -1,92 | 12 | -2,14 | 12 | -2,11 | 12 | -2,25 |
| | 13 | -1,91 | 13 | -2,13 | 13 | -2,11 | 13 | -2,24 |
| | 9 | -3,16 | 9 | -3,38 | 9 | -3,32 | 9 | -3,31 |



S5-G5

| | | | | | | | | |
|---|---|---|---|---|---|---|---|---|
| | 7 | -2,54 | 7 | -2,71 | 7 | -2,77 | 7 | -2,72 |
| | 4 | -2,39 | 4 | -2,52 | 4 | -2,59 | 4 | -2,54 |
| | 8 | -2,28 | 8 | -2,44 | 8 | -2,53 | 8 | -2,46 |
| | 5 | -2,04 | 3 | -2,17 | 3 | -2,24 | 5 | -2,18 |
| | 3 | -2,04 | 5 | -2,16 | 5 | -2,23 | 3 | -2,18 |
| | 11 | -1,93 | 10 | -2,09 | 12 | -2,14 | 10 | -2,10 |
| | 12 | -1,93 | 11 | -2,08 | 11 | -2,14 | 12 | -2,09 |
| | 10 | -1,93 | 12 | -2,08 | 10 | -2,13 | 11 | -2,08 |
| | 6 | -1,89 | 6 | -2,03 | 6 | -2,09 | 6 | -2,03 |
| | 2 | -1,89 | 2 | -2,02 | 2 | -2,08 | 2 | -2,03 |
| | 1 | -1,85 | 1 | -1,98 | 1 | -2,04 | 1 | -1,99 |
| S5-G6 | 4 | -2,45 | 4 | -2,59 | 4 | -2,62 | 4 | -2,64 |
| | 7 | -2,38 | 10 | -2,57 | 10 | -2,58 | 10 | -2,59 |
| | 10 | -2,37 | 7 | -2,51 | 7 | -2,56 | 7 | -2,58 |
| | 5 | -2,22 | 5 | -2,35 | 5 | -2,39 | 5 | -2,39 |
| | 3 | -2,18 | 3 | -2,32 | 3 | -2,36 | 3 | -2,39 |
| | 8 | -2,08 | 8 | -2,23 | 8 | -2,27 | 8 | -2,31 |
| | 6 | -2,07 | 6 | -2,20 | 6 | -2,24 | 6 | -2,25 |
| | 9 | -2,02 | 9 | -2,16 | 9 | -2,19 | 9 | -2,21 |
| | 11 | -1,71 | 11 | -1,91 | 11 | -1,96 | 11 | -1,98 |
| | 12 | -1,71 | 12 | -1,90 | 12 | -1,95 | 12 | -1,98 |
| | 1 | -1,34 | 1 | -1,47 | 1 | -1,48 | 1 | -1,48 |
| | 2 | -1,31 | 2 | -1,43 | 2 | -1,45 | 2 | -1,46 |

Table S3: Fingerprint values for all the SAMPL5 ligands for different water models. The atom numbering refers to the ligand heavy atoms as numbered in the PDB input files.

| Ligand | TIP3P | | SPC | | TIP4P | | TIP4Pew | |
|---|---|---|---|---|---|---|---|---|
| | Atom | Fingerprint | Atom | Fingerprint | Atom | Fingerprint | Atom | Fingerprint |
| S6-G0 | 6 | -3,13 | 6 | -3,26 | 6 | -3,27 | 6 | -3,36 |
| | 1 | -2,58 | 1 | -2,80 | 1 | -2,78 | 1 | -2,85 |
| | 5 | -2,51 | 5 | -2,63 | 5 | -2,64 | 5 | -2,68 |
| | 4 | -2,50 | 4 | -2,61 | 4 | -2,62 | 7 | -2,67 |
| | 7 | -2,47 | 7 | -2,59 | 7 | -2,61 | 4 | -2,66 |
| | 3 | -2,40 | 3 | -2,52 | 3 | -2,54 | 3 | -2,59 |
| | 2 | -2,39 | 2 | -2,51 | 2 | -2,53 | 2 | -2,59 |
| | 8 | -1,84 | 9 | -2,00 | 8 | -2,01 | 8 | -2,11 |
| | 9 | -1,83 | 8 | -2,00 | 9 | -1,99 | 9 | -2,09 |
| S6-G1 | 3 | -2,59 | 3 | -2,76 | 3 | -2,76 | 6 | -2,77 |
| | 6 | -2,57 | 6 | -2,70 | 6 | -2,76 | 3 | -2,75 |
| | 5 | -2,43 | 5 | -2,54 | 5 | -2,59 | 5 | -2,60 |
| | 4 | -2,27 | 4 | -2,40 | 4 | -2,47 | 4 | -2,49 |
| | 2 | -2,06 | 2 | -2,17 | 2 | -2,20 | 2 | -2,21 |
| | 1 | -1,99 | 1 | -2,11 | 1 | -2,17 | 1 | -2,18 |
| | 7 | -1,77 | 8 | -1,96 | 8 | -2,02 | 7 | -2,07 |
| | 8 | -1,77 | 7 | -1,95 | 7 | -2,00 | 8 | -2,06 |
| S6-G2 | 9 | -3,44 | 9 | -3,55 | 9 | -3,63 | 9 | -3,58 |
| | 5 | -2,95 | 5 | -3,09 | 5 | -3,17 | 5 | -3,13 |
| | 8 | -2,73 | 8 | -2,85 | 8 | -2,92 | 6 | -2,85 |
| | 6 | -2,69 | 6 | -2,81 | 6 | -2,90 | 8 | -2,85 |
| | 4 | -2,52 | 4 | -2,71 | 4 | -2,73 | 4 | -2,67 |
| | 2 | -2,39 | 2 | -2,53 | 2 | -2,63 | 2 | -2,59 |
| | 7 | -2,37 | 7 | -2,51 | 7 | -2,58 | 7 | -2,52 |
| | 10 | -2,31 | 10 | -2,46 | 10 | -2,55 | 10 | -2,49 |

| | | | | | | | | |
|---|---|---|---|---|---|---|---|---|
| | 1 | -2,10 | 1 | -2,24 | 1 | -2,31 | 1 | -2,25 |
| | 3 | -1,88 | 3 | -1,99 | 3 | -2,06 | 12 | -2,04 |
| | 11 | -1,80 | 11 | -1,98 | 12 | -2,04 | 11 | -2,03 |
| | 12 | -1,79 | 12 | -1,97 | 11 | -2,03 | 3 | -1,99 |
| S6-G3 | 3 | -2,54 | 3 | -2,77 | 3 | -2,73 | 3 | -2,79 |
| | 5 | -2,50 | 5 | -2,65 | 5 | -2,66 | 5 | -2,74 |
| | 6 | -2,50 | 6 | -2,63 | 6 | -2,64 | 6 | -2,70 |
| | 4 | -2,40 | 4 | -2,53 | 4 | -2,53 | 4 | -2,59 |
| | 2 | -1,99 | 2 | -2,13 | 2 | -2,12 | 2 | -2,19 |
| | 7 | -1,80 | 7 | -1,99 | 7 | -1,98 | 7 | -2,08 |
| | 8 | -1,80 | 8 | -1,98 | 8 | -1,97 | 8 | -2,06 |
| | 1 | -1,63 | 1 | -1,76 | 1 | -1,74 | 1 | -1,81 |
| S6-G4 | 10 | -3,34 | 10 | -3,52 | 10 | -3,52 | 10 | -3,62 |
| | 2 | -2,79 | 2 | -3,00 | 2 | -2,99 | 2 | -3,04 |
| | 9 | -2,74 | 3 | -2,92 | 3 | -2,91 | 3 | -2,97 |
| | 7 | -2,73 | 9 | -2,89 | 7 | -2,89 | 9 | -2,89 |
| | 3 | -2,69 | 7 | -2,88 | 9 | -2,88 | 7 | -2,88 |
| | 8 | -2,49 | 8 | -2,66 | 8 | -2,67 | 8 | -2,70 |
| | 5 | -2,35 | 5 | -2,56 | 5 | -2,57 | 4 | -2,60 |
| | 4 | -2,30 | 4 | -2,53 | 4 | -2,55 | 5 | -2,60 |
| | 6 | -2,30 | 6 | -2,50 | 6 | -2,53 | 6 | -2,54 |
| | 1 | -2,26 | 1 | -2,40 | 1 | -2,37 | 1 | -2,35 |
| | 11 | -1,92 | 11 | -2,12 | 11 | -2,11 | 12 | -2,20 |
| | 12 | -1,91 | 12 | -2,11 | 12 | -2,11 | 11 | -2,19 |
| S6-G5 | 6 | -2,92 | 6 | -3,10 | 6 | -3,11 | 6 | -3,16 |
| | 3 | -2,73 | 3 | -3,06 | 3 | -2,96 | 3 | -3,04 |
| | 5 | -2,42 | 5 | -2,59 | 5 | -2,58 | 5 | -2,60 |
| | 4 | -2,27 | 4 | -2,49 | 4 | -2,48 | 4 | -2,53 |
| | 2 | -1,90 | 8 | -2,11 | 7 | -2,08 | 7 | -2,17 |
| | 7 | -1,85 | 7 | -2,10 | 8 | -2,06 | 8 | -2,15 |
| | 8 | -1,83 | 2 | -2,08 | 2 | -2,06 | 2 | -2,07 |
| | 1 | -1,59 | 1 | -1,77 | 1 | -1,73 | 1 | -1,76 |
| S6-G6 | 6 | -3,33 | 6 | -3,49 | 6 | -3,56 | 6 | -3,64 |
| | 5 | -2,61 | 1 | -2,81 | 1 | -2,82 | 1 | -2,86 |
| | 1 | -2,59 | 5 | -2,72 | 5 | -2,75 | 5 | -2,76 |
| | 4 | -2,52 | 4 | -2,64 | 4 | -2,69 | 4 | -2,71 |
| | 2 | -2,28 | 2 | -2,41 | 2 | -2,46 | 2 | -2,49 |
| | 3 | -2,26 | 3 | -2,40 | 3 | -2,44 | 3 | -2,47 |
| | 7 | -1,83 | 7 | -2,01 | 7 | -2,03 | 7 | -2,09 |
| | 8 | -1,81 | 8 | -2,00 | 8 | -2,00 | 8 | -2,07 |
| S6-G7 | 7 | -4,10 | 7 | -4,21 | 7 | -4,30 | 7 | -4,44 |
| | 3 | -2,97 | 3 | -3,22 | 3 | -3,09 | 3 | -3,17 |
| | 6 | -2,67 | 6 | -2,79 | 6 | -2,78 | 6 | -2,84 |
| | 4 | -2,39 | 4 | -2,50 | 5 | -2,46 | 4 | -2,62 |
| | 5 | -2,30 | 5 | -2,45 | 4 | -2,45 | 5 | -2,53 |
| | 2 | -2,04 | 2 | -2,16 | 2 | -2,15 | 2 | -2,23 |
| | 8 | -1,97 | 8 | -2,14 | 8 | -2,10 | 8 | -2,22 |
| | 9 | -1,97 | 9 | -2,12 | 9 | -2,10 | 9 | -2,22 |
| | 1 | -1,75 | 1 | -1,89 | 1 | -1,86 | 1 | -1,96 |

Table S4: Fingerprint values for all the SAMPL6 ligands for different water models. The atom numbering refers to the ligand heavy atoms as numbered in the PDB input files.

### Example of PLUMED input file for an OPES simulation

To ensure reproducibility, we explicitly report an example of PLUMED input file to perform the OPES simulations of the host–guest complexes. The file specifies the chosen CVs, biasing parameters, and simulation settings. All the files used for the simulations are collected in the Github repository reported in the main text.

```
# -- (1) ATOMS DEFINITIONS and ALIGNMENT --


HOST: GROUP ATOMS=18-213 #host atoms
LIGC: GROUP ATOMS=1-8 #carbon atoms in the ligand
l1:  GROUP ATOMS=3 #ligand selected atoms
l2:  GROUP ATOMS=5
l3:  GROUP ATOMS=5
l4:  GROUP ATOMS=8
WO: GROUP ATOMS=214-6831:3 #water oxygen atoms
WHOLEMOLECULES ENTITY0=HOST
FIT_TO_TEMPLATE STRIDE=1 REFERENCE=conf_template.pdb
TYPE=OPTIMAL #coordinates alignment
lig:  CENTER ATOMS=LIGC


v1:  FIXEDATOM AT=2.0136,2.0136,2.0 #virtual atoms
v2:  FIXEDATOM AT=2.0136,2.0136,2.25
v3:  FIXEDATOM AT=2.0136,2.0136,2.5
v4:  FIXEDATOM AT=2.0136,2.0136,2.75
v5:  FIXEDATOM AT=2.0136,2.0136,3.0
v6:  FIXEDATOM AT=2.0136,2.0136,3.25
v7:  FIXEDATOM AT=2.0136,2.0136,3.5
v8:  FIXEDATOM AT=2.0136,2.0136,3.75


cyl:  DISTANCE ATOMS=v1,lig COMPONENTS
radius:  MATHEVAL ARG=cyl.x,cyl.y FUNC=sqrt(x*x+y*y)
PERIODIC=NO


# -- (2) DESCRIPTORS --


L1:  COORDINATION GROUPA=l1 GROUPB=WO SWITCH=RATIONAL D_0=0.0 R_0=0.35 NN=6 MM=10 D_MAX=1.20
NLIST NL_CUTOFF=1.6 NL_STRIDE=20 #in nm
L2:  COORDINATION GROUPA=l2 GROUPB=WO SWITCH=RATIONAL D_0=0.0 R_0=0.35 NN=6 MM=10 D_MAX=1.20
NLIST NL_CUTOFF=1.6 NL_STRIDE=20
V2:  COORDINATION GROUPA=v2 GROUPB=WO SWITCH=RATIONAL D_0=0.0 R_0=0.35 NN=2 MM=6 D_MAX=1.20 NLIST
NL_CUTOFF=1.6 NL_STRIDE=20

L3:  COORDINATION GROUPA=l3 GROUPB=WO SWITCH=RATIONAL D_0=0.0 _0=0.25 NN=6 MM=10 D_MAX=0.8 NLIST
NL_CUTOFF=1.5 NL_STRIDE=20
L4:  COORDINATION GROUPA=l4 GROUPB=WO SWITCH=RATIONAL D_0=0.0 R_0=0.25 NN=6 MM=10 D_MAX=0.8 NLIST
NL_CUTOFF=1.5 NL_STRIDE=20
V1:  COORDINATION GROUPA=v1 GROUPB=WO SWITCH=RATIONAL D_0=0.0 R_0=0.25 NN=2 MM=6 D_MAX=0.8 NLIST
NL_CUTOFF=1.5 NL_STRIDE=20
V3:  COORDINATION GROUPA=v3 GROUPB=WO SWITCH=RATIONAL D_0=0.0 R_0=0.25 NN=2 MM=6 D_MAX=0.8 NLIST
NL_CUTOFF=1.5 NL_STRIDE=20
V4:  COORDINATION GROUPA=v4 GROUPB=WO SWITCH=RATIONAL D_0=0.0 R_0=0.25 NN=2 MM=6 D_MAX=0.8 NLIST
NL_CUTOFF=1.5 NL_STRIDE=20
V5:  COORDINATION GROUPA=v5 GROUPB=WO SWITCH=RATIONAL D_0=0.0 R_0=0.25 NN=2 MM=6 D_MAX=0.8 NLIST
NL_CUTOFF=1.5 NL_STRIDE=20
V6:  COORDINATION GROUPA=v6 GROUPB=WO SWITCH=RATIONAL D_0=0.0 R_0=0.25 NN=2 MM=6 D_MAX=0.8 NLIST
NL_CUTOFF=1.5 NL_STRIDE=20
V7:  COORDINATION GROUPA=v7 GROUPB=WO SWITCH=RATIONAL D_0=0.0 R_0=0.25 NN=2 MM=6 D_MAX=0.8 NLIST
NL_CUTOFF=1.5 NL_STRIDE=20
```

```
V8:  COORDINATION GROUPA=v8 GROUPB=WO SWITCH=RATIONAL D_0=0.0 R_0=0.25 NN=2 MM=6 D_MAX=0.8 NLIST
NL_CUTOFF=1.5 NL_STRIDE=20

# -- (3) Funnel, walls and angle definitions --


ang:  ANGLE ATOMS=v3,v5,1,8 #angle of a ligand's axis with z
cosang:  MATHEVAL ARG=ang FUNC=cos(x) PERIODIC=NO


funnel:  MATHEVAL ARG=radius,cyl.z VAR=r,z FUNC=(r+1.0*(-1.2+z))*step(-z+1.)+(r-0.2)*step(z-1.)
PERIODIC=NO
UPPER_WALLS AT=0 ARG=funnel KAPPA=2000.0 LABEL=funnelwall #funnel restraint
UPPER_WALLS AT=1.8 ARG=cyl.z KAPPA=4000.0 EXP=2 LABEL=upper_wall #upper limit of cyl.z


ene:  ENERGY


# -- (4) OPES --
  OPES_METAD_EXPLORE ...
LABEL=opes
ARG=L1,L2,V2
#SIGMA=0.077,0.48
FILE=Kernels.data
STATE_RFILE=compressed.Kernels.data
STATE_WFILE=compressed.Kernels.data
PACE=10000
BARRIER=50
...  OPES_METAD_EXPLORE


PRINT
ARG=opes.bias,cyl.z,radius,funnelwall.bias,upper_wall.bias,ene,cosang,L1,L2,L3,L4,V1,V2,V3,V4,V5,V6,V7,V8
STRIDE=500 FILE=COLVAR FMT=%8.4f
```
